# Proviruses in CD4^+^ T cells reactive to autologous antigens contribute to nonsuppressible HIV-1 viremia

**DOI:** 10.1101/2025.04.19.649587

**Authors:** Fengting Wu, Milica Moskovljevic, Filippo Dragoni, Sahana Jayaraman, Nathan L. Board, Angelica Camilo-Contreras, Silvia Bernal, Vivek Hariharan, Hao Zhang, Jun Lai, Anushka Singhal, Sebastien Poulin, Frederic Chano, Annie Chamberland, Cecile Tremblay, Meredith Zoltick, Christopher J. Hoffmann, Joyce L. Jones, H. Benjamin Larman, Luis J. Montaner, Janet D. Siliciano, Robert F. Siliciano, Francesco R. Simonetti

**Affiliations:** Department of Medicine, Johns Hopkins University School of Medicine, Baltimore, MD, USA; Institute of Cell Engineering, Division of Immunology, Department of Pathology, Johns Hopkins University, Baltimore, MD, USA; Department of Molecular Microbiology and Immunology, Johns Hopkins Bloomberg School of Public Health, Baltimore, MD, USA; HIV Cure and Viral Diseases Center, The Wistar Institute, Philadelphia, PA, USA; Howard Hughes Medical Institute, Baltimore, MD, USA; Clinique L’Agora, Montréal, Québec, Canada; Centre de Recherche du Centre Hospitalier de l’Université de Montréal, Montréal, Québec, Canada; Département de Microbiologie, Immunologie et Infectiologie, Université de Montréal, Montréal, Québec, Canada

## Abstract

Antiretroviral therapy (ART) halts HIV-1 replication, reducing plasma virus levels to below the limit of detection, but it is not curative due to a reservoir of latently infected CD4^+^ T cells. In some people living with HIV-1 (PLWH), plasma HIV-1 RNA becomes persistently detectable despite optimal ART. This nonsuppressible viremia (NSV) is characterized by identical, non-evolving HIV-1 RNA variants expressed from infected CD4^+^ T cell clones. The mechanisms driving persistent virus production from a specific population of infected cells are poorly understood. We hypothesized that proviruses in cells responding to chronic immunologic stimuli, including self-associated antigens, may drive viral gene expression and NSV. Here, we demonstrate that stimulation of CD4^+^ T cells with autologous cell lysates induces virus production in an MHC-II-dependent manner. In 7 of 8 participants with NSV, we recovered viral RNA released ex vivo in response to autologous cell lysates that matched plasma virus. This process involves both defective and replication-competent proviruses residing in conventional T cells, and is also observed in PLWH with undetectable viremia. These findings suggest that recognition of self-associated antigens is an important cause of HIV-1 reservoir expression, which can contribute to persistent systemic inflammation and potential rebound upon ART interruption.

**One sentence summary:** HIV-1 viremia not suppressed by effective ART can be caused by proviruses in CD4^+^ T cells reactive to autologous antigens.

## INTRODUCTION

The latent reservoir poses a formidable challenge to curing HIV-1 infection. Although antiretroviral therapy (ART) effectively halts HIV-1 replication and decreases plasma viral load (VL) to below the detection limit of clinical assays, it is not curative (*1–3*). A small population of infected memory CD4^+^ T cells carrying latent HIV-1 genomes persists for life in people living with HIV-1 (PLWH) despite decades of effective ART (*4–8*). These latently infected cells can evade immune surveillance and clonally expand in response to antigens and other immunologic stimuli, thus maintaining the latent reservoir (*9–14*).

Even with 100% adherence to ART, a large fraction of PLWH have residual viremia (RV, typically 1-3 copies/mL) detectable by research-based ultrasensitive assays (*15–18*). In these cases, intensifying or optimizing ART fails to reduce RV(*19, 20*). Sequencing of RV has revealed clonal populations that lack drug resistance mutations and do not evolve over time (*21–23*). Furthermore, a rare group of PLWH experience higher levels of RV in the detectable range (VL >20-50 copies/mL) that can persist for years. This clinical scenario is often referred to as nonsuppressible viremia (NSV), to highlight the inability of ART optimization or intensification to further reduce viremia below the limit of detection. Although most cases of detectable viremia would fall within the WHO and UNAIDS definition of suppression for the elimination of viral transmission (<200cp/mL), NSV remains a cause of distress for PLWH and their care providers. Furthermore, clinical options for managing NSV are currently limited (*24, 25*). Halvas *et al.* characterized plasma virus from a small cohort of people with NSV and showed that some predominant plasma clones (PPC) were identical to sequences of virus recovered from viral outgrowth assays, suggesting that NSV is caused by clonally expanded T cells carrying replication-competent proviruses (*26*). However, White *et al*. recently demonstrated that replication incompetent proviruses with 5’-Leader defects, affecting the major splicing donor (MSD) site, are also a common cause of NSV. Small deletions in the MSD necessitate inefficient alternative splicing, which results in decreased levels of HIV-1 Envelope incorporation into the virion, thereby abrogating replication competence (*27*). Although different in magnitude, RV and NSV are manifestations of the same phenomenon, reflecting the pervasive reservoir persistence via cell proliferation and expression of viral genes from a subset of infected cells, which occurs in all PLWH(*26–30*). However, the virological or cellular factors driving the expression of proviruses that contribute to viremia on ART and its consequences remain unclear. Moreover, proviruses with spontaneous transcriptional activity during ART may be the first to contribute to viral rebound upon treatment interruption (*31–33*). Therefore, understanding the causes of persistent viremia will be critical for improving HIV-1 clinical care and developing curative strategies.

RV and NSV are usually characterized by one or a few clonal variants in plasma, suggesting that they are maintained by the ongoing stimulation of specific T cells. The engagement of T cell receptors (TCR) with peptides presented by major histocompatibility complex class II (MHC-II) molecules on antigen-presenting cells (APCs) is essential for initiating CD4^+^ T cell activation, effector function, proliferation, and maintenance of immunological memory (*34–36*). Additionally, recent studies from our group and others have shown that an encounter with cognate antigens can lead to the induction of HIV-1 gene expression, demonstrating a direct relationship between antigen recognition and HIV-1 latency reversal (*37–41*). MHC-II restricted epitopes can be derived from both exogenous and endogenous sources, ranging from foreign pathogens or apoptotic cells in the microenvironment to intracellular proteins within APCs (*42–44*). Moreover, CD4^+^ T cell responses can also exhibit cross-reactivity (also known as heterologous immunity), wherein a TCR can recognize two or more distinct peptide:MHC-II (pMHC-II) complexes (*45, 46*). Recognition of self-antigens is essential for the persistence and homeostasis of naïve and memory cells (*47–49*). However, recognition of self:pMHC is tightly regulated, as self-tolerance loss can result in autoimmunity (*48, 50*). Whether engagement with self-associated pMHC can reverse HIV-1 latency and contribute to spontaneous reservoir activity is unknown.

Previous studies have demonstrated the role of various foreign antigens in driving the persistence and dynamics of HIV-1-infected CD4^+^ T cell clones (*12, 13, 51–53*). We hypothesized that transcriptionally active proviruses, including those causing persistent viremia, may be present in cells reactive to chronic antigenic stimuli, such as autologous peptides that are constitutively presented by APCs. Here, we provide evidence that recognition of autologous cell lysates by CD4^+^ T cells ex vivo can induce latency reversal and production of virions identical to those found in the plasma of PLWH on ART.

## RESULTS

### NSV is driven by oligoclonal HIV-1 variants

We characterized eight individuals with persistent NSV, defined as plasma HIV-1 RNA above 20 copies/mL for more than 12 months on optimal ART. Study participants were referred to us by care providers due to NSV despite optimal adherence to ART and no history of drug resistance. Their characteristics and clinical history are summarized in Figure 1A and Supplementary Figure S1. All participants were on long-term suppressive ART (median 27 years, range 9-31 years) until they developed persistent NSV for a median of 5 years (range 3-10 years). Plasma HIV-1 RNA levels varied, with a median of 45 copies/mL (range 31.6-3400 copies/mL). Single genome sequencing in the P6-RT region demonstrated that plasma viruses belonged to subtype B and lacked drug resistance mutations. As observed previously, the plasma virus was dominated by sets of identical sequences despite extensive pre-ART viral diversification (*23*). Some participants had multiple plasma clones (P2, P5, P6, P7, and P8), while other participants had a single dominant clone (P1, P3, and P4) contributing to NSV (Figure 1B). Furthermore, we recently demonstrated that proviruses with 5’-Leader defects contributed to NSV in P1, P2, P3, and P4 (*27*). A large fraction of sequences found in P6 and P7 were identical to the virus recovered from p24^+^ wells of the quantitative viral outgrowth assay (QVOA), indicating that their PPCs were replication-competent (Figure 1B). Although sequencing of the P6-RT region in P5 and P8 showed no significant defects, we found no matching replication-competent QVOA isolates; therefore, their PPCs may have defects in other regions of the genome. These results confirm that NSV can originate from replication-competent proviruses and proviruses with 5’-Leader defects.

**Figure 1.**
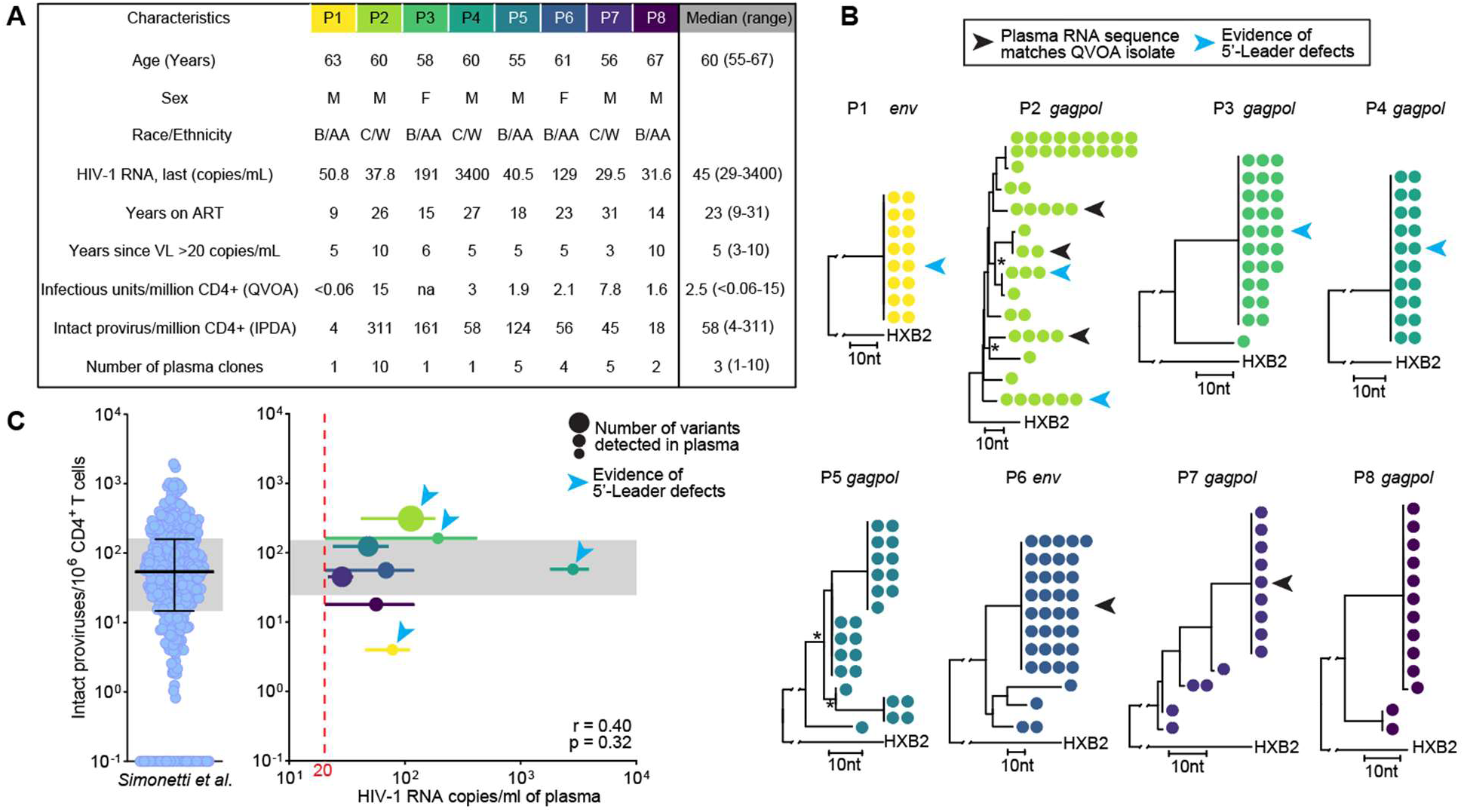
Clinical characteristics, HIV sequences in plasma, and reservoir size participants with NSV. **(A)** Clinical characteristics and reservoir size measurements; “na” indicates not available. **(B)** Analysis of HIV-1 RNA plasma sequences; maximum likelihood trees are rooted on the HXB2 reference; variants matching sequences recovered from the quantitative viral outgrowth assay (QVOA) are indicated by black arrowheads; variants with known defects in the 5’-Leader are indicated with blue arrowheads. **(C)** Comparison of intact proviral DNA frequencies by IPDA from 400 people on ART with undetectable viremia(*54*)(left) and the participants in this study (right); circles in the right panel match colors as in A and B; horizontal bars represent the range of plasma HIV-1 RNA during the period of persistent viremia, and circles indicate median values; circle size indicates the number of variants found in plasma; the red dashed line represents the limit of the detection of clinical assays of 20 copies/mL, and the gray area indicates the interquartile range of the frequency of intact proviruses.

Analysis of HIV-1 reservoir size in study participants showed a median of 2.55 infectious units per million (IUPM, range 0.06-15) by the QVOA, and a median frequency of 58 intact HIV-1 proviruses/10^6^ CD4^+^ T cells (range 4-311) by intact proviral DNA assay (IPDA) (Figure 1A and C). The median frequency of intact proviruses by IPDA in NSV participants did not differ from that of 400 individuals on ART with undetectable HIV-1 RNA, as previously reported (*54*). We observed no correlation between reservoir size and level of viral load for our study participants (r=0.40, p=0.32) (Figure 1C), suggesting that reservoir size is not sufficient to explain the development of NSV (*16, 55*).

### Autologous cell lysates induce virus production in an MHC-II-dependent manner

To test the hypothesis that recognition of self-associated antigens can induce virion production from latently infected cells, we isolated CD4^+^ T cells from study participants and cultured them with autologous monocyte-derived dendritic cells (DCs) loaded with lysates of autologous cells from various lineages. DCs were generated from autologous CD14^+^ monocytes by incubation with IL-4 and GM-CSF for 5 days, generating DCs capable of efficiently presenting exogenous antigens (*56*). Autologous lysates were generated by sorting peripheral blood mononuclear cells (PBMCs) based on lineage markers to isolate B cells (CD19^+^ or CD20^+^CD3^-^), T cells (CD19^-^CD20^-^ CD3^+^), and non-B-non-T cells (nBnT, CD19^-^CD20^-^CD3^-^) (Figure 2A, Supplementary Figures S2A and S2B, and see Methods). These purified cell populations were then lysed by multiple freeze-thaw cycles, and insoluble components were removed by centrifugation. The distribution of immune cell lineages varied across participants, with a mean of 9.4% B cells, 52.1% T cells, and 35.8% nBnT (Supplementary Figure S2C). CD4^+^ T cells were cultured with antigen-pulsed DCs for 7 days in the presence of antiretroviral drugs. As negative and positive controls, respectively, we cultured CD4^+^ T cells without stimulation and CD4+ T cells nonspecifically stimulated by ɑCD3/CD28 antibodies (see Methods). Supernatants were collected on days 1, 2, 3, 5, and 7 of culture. To test whether infected T-cells produced virions in response to recognition of specific self-associated antigens presented by DCs, we included a control with a pan-MHC-II blocking antibody, Tü39, for each experimental condition. Viral RNA was extracted from virions pelleted by centrifugation, quantified by digital PCR (dPCR) assays targeting the polyadenylated 3’ end of HIV-1 RNA (polyA assay) or provirus-specific regions, and sequenced by SGS (*57, 58*) (Figure 2B). Finally, to exclude that virus production could derive from the autologous DCs in the coculture (*59*), we quantified total and intact HIV-1 DNA in CD14^+^ monocytes from four participants. Across all samples, we detected only a single copy of HIV-1 LTR in P6, corresponding to >400-fold lower infection frequency in monocytes than in CD4^+^ T cells (Supplementary Figure S2 D-F), strongly suggesting that monocytes are not the source of virus in our culture model.

**Figure 2.**
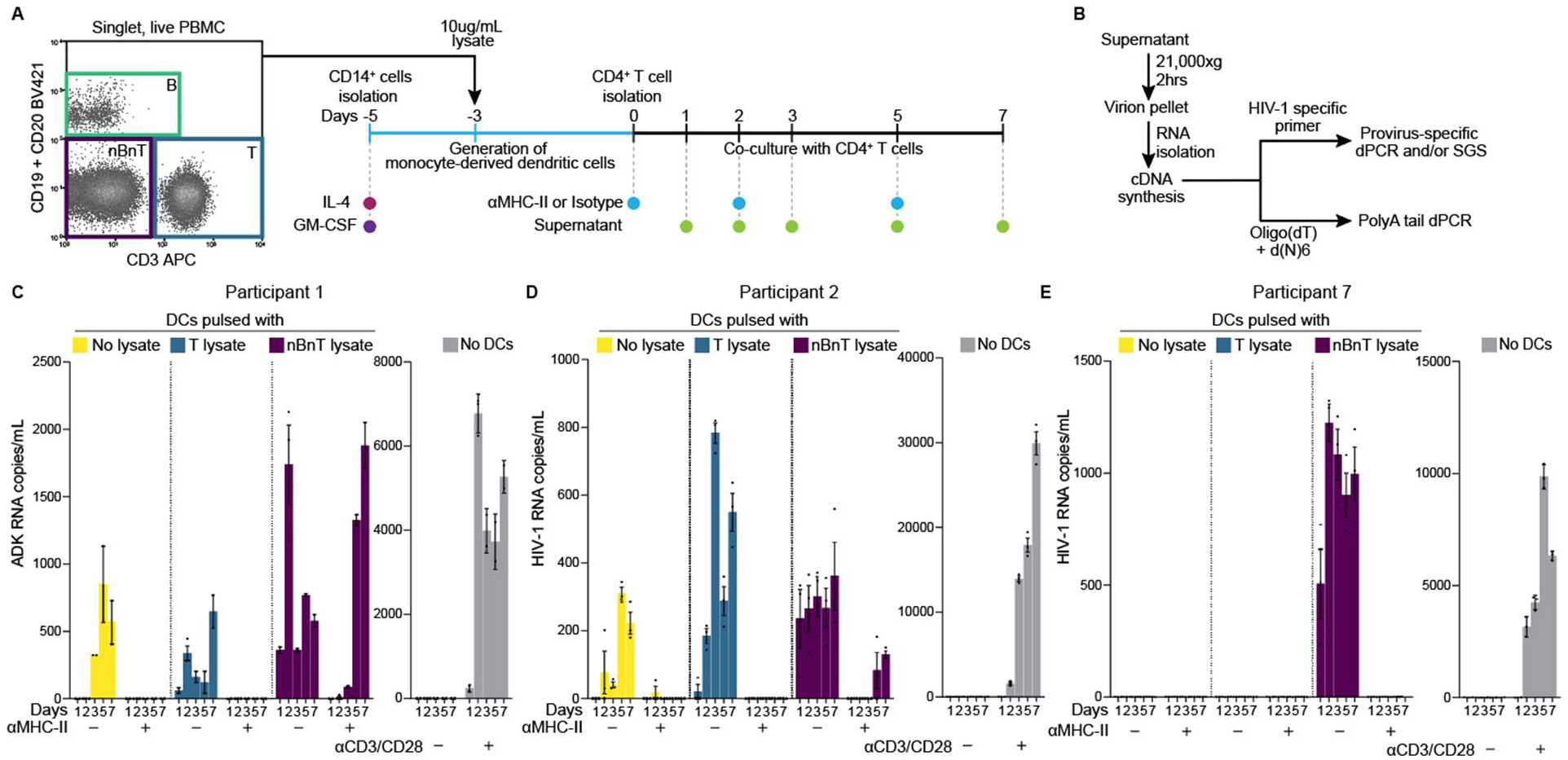
Experimental approach to investigate autologous lysate-induced virus production ex vivo. **(A)** Experimental design and flow gating strategy to isolate cells and generate autologous lysates. Participant’s PBMCs were sorted based on viability, CD19, CD20, and CD3, followed by lysis through freeze-thaw cycles and protein quantification. DCs were generated as previously described(*56*), and coculture was set up with CD4^+^ T cells from each participant. **(B)** Workflow to isolate HIV-1 RNA from virions from culture supernatant and cDNA synthesis for downstream assays. **(C)** Representative plots of HIV-1 RNA measurements from the ADK.d22 provirus for participant P1. CD4^+^ T cells were cultured with and without DCs. **(D and E)** Representative plots of HIV-1 PolyA RNA values for participants P2 and P7. Conditions with ɑMHC-II antibody or isotype control are indicated below the x-axis. Dots represent dPCR replicate measurements, while error bars indicate the mean and standard error.

Figures 2C-E show the results from three representative participants with NSV, P1, P2, and P7. For P1, virus production in supernatant was detected with an assay specific to the ADK.d22 provirus to target the main viral variant detected in plasma for this participant (*27*). In P1, we observed virus production in response to DCs pulsed with autologous T and nBnT cell lysates. Interestingly, some virus production was also detected when CD4^+^ T cells were cultured with unpulsed DCs (Figure 2C). This observation may reflect that DCs alone can process and present endogenous intracellular proteins or exogenous antigens taken from surrounding apoptotic cells and extracellular vesicles (*42–44*). Similarly, in P2 we observed virus production in response to unpulsed DCs and DCs pulsed with autologous T and nBnT cell lysates using the HIV-1 polyA assay (Figure 2D). In P7, virus production by CD4^+^ T cells was only observed when cultured with nBnT-lysate pulsed DCs (Figure 2E). Overall, we detected virus production by CD4^+^ T cells cocultured with unpulsed DCs in 5 of 8 NSV participants (see below). The addition of the ɑMHC-II antibody either completely blocked, significantly decreased, or delayed virus production compared to the isotype control (Figure 2C-E). Despite these results, we saw virus production in some culture wells treated with the Tü39 clone, potentially due to reduced efficiency in binding to specific HLA proteins or incomplete blockade of TCR-pMHC interactions (*60*). To validate the effect of the ɑMHC-II antibody to block MHC-II-mediated responses, we stimulated CD8-depleted PBMCs from P1 with CMV lysate for 18 hours. The presence of ɑMHC-II antibody did not affect cell viability for CD4^+^ T cells. The addition of the ɑMHC-II blocking antibody reduced the percentages of CD154^+^CD69^+^ and CD154^+^CD137^+^ in CD3^+^CD4^+^ T cells by 2.6-fold in both cases (Supplementary Figure S3A and S3B). In addition, the proliferation of CMV-responding CD3^+^CD4^+^ T cells after 7 days was markedly decreased by ɑMHC-II blockade compared to isotype control (Supplementary Figure S3C), demonstrating that the addition of ɑMHC-II blocking antibody decreases both cell activation and proliferation in response to antigenic stimulation (*61*). We analyzed the impact of the ɑMHC-II blocking antibody on virus production parsed by participants to investigate whether specific MHC-II alleles could affect the ɑMHC-blockade efficiency, but we observed no apparent differences (Supplementary Figure S3D).

To estimate overall virus production, we calculated the area under the curve (AUCg) of HIV-1 RNA measured over the 7 days of coculture (*62*). Using the HIV-1 RNA AUCg as a metric helps mitigate the bias introduced by removing and replacing 200µL culture media on days 1, 2, 3, 5, and 7. Of the 8 NSV participants studied, 7 had detectable virus release by CD4^+^ T cells in response to stimulation with lysate-pulsed DCs, in some cases with one or more autologous lysates (Figure 3A). All 7 participants showed virus production by CD4^+^ T cells in response to nBnT lysate. Additionally, 5 out of 7 participants had CD4^+^ T cells that produced virus in response to unpulsed DCs. CD4^+^ T cells from three participants (P1, P2, P3) also exhibited additional virus release when stimulated with either B or T cell lysates (Figure 3A). Unstimulated CD4^+^ T cells produced little or no virus across all participants, suggesting that spontaneous latency reversal from highly inducible proviruses cannot explain our results (Figure 3B). Stimulation with autologous lysates induced 24.7-fold higher HIV-1 RNA expression than unstimulated CD4^+^ T cells (p=0.035), indicating a response against antigens presented by DCs. CD4^+^ T cells stimulated with ɑCD3/CD28 resulted in even higher virus production than coculture with DCs (37.4-fold, p=0.01) and CD4^+^ T cells left untreated (921.2-fold, p<0.0001), reflecting nonspecific polyclonal activation of CD4^+^ T cells (Figure 3B and 3C). Overall, we observed significantly reduced virus production (p=0.03) with the ɑMHC-II block (Figure 3D and Supplementary Figure S3D).

**Figure 3.**
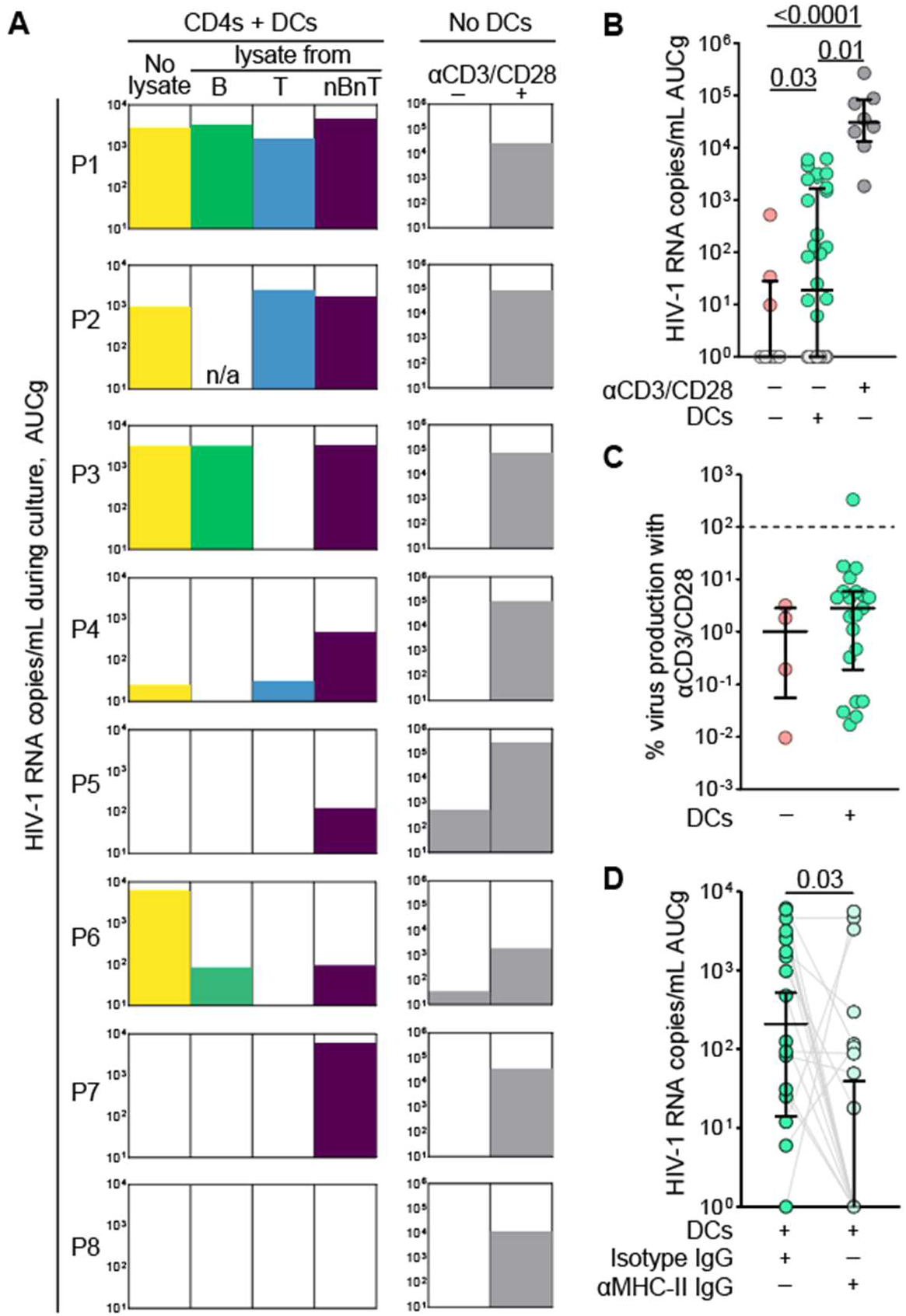
Autologous lysates induce virion production in an MHC-II-dependent manner. **(A)** Area under the curve of HIV-1 PolyA RNA across 7 days of culture for all participants with NSV; CD4+ T cell culture conditions are indicated at the top of the graph; "n/a" represents conditions that were not tested due to insufficient amount of cell lysate. **(B)** HIV-1 PolyA RNA AUCg values across all experiments grouped by type of stimulation; symbol colors indicate untreated CD4s (pink), coculture of CD4s with pulsed or unpulsed DCs (green), and CD4s treated with ɑCD3/CD28 (grey). **(C)** AUCg values obtained from CD4s with and without DCs were used to calculate the percentage of virus production compared to stimulation with ɑCD3/CD28. **(D)** AUCg values of virus production upon stimulation with autologous lysates from all participants using isotype control versus ɑMHC-II blocking antibody. Statistical significance was calculated by a two-tailed paired t-test.

Foreign antigens contribute to clonal expansion of infected cells in vivo (*12, 13, 63*) and latency reversal ex vivo (*40, 41*). Thus, we also examined virus production by CD4^+^ T cells incubated with autologous DCs pulsed HIV-1 Gag p55. We selected two participants known to have HIV-1-infected, Gag responding cells (Supplementary Figure S4). Stimulation of CD4^+^ T cells with HIV-1 Gag pulsed DCs resulted in virus production in both participants (P2 and P10). Single genome sequences of virus-associated HIV-1 RNA in supernatant matched the proviruses recovered from Gag-responding cells isolated in previous experiments (*41*). In P2, stimulation with Gag-pulsed DCs resulted in the production of virus identical to a clone found in plasma, which displays a defect in the MSD and is integrated into the *RRM1* gene (*27*). Conversely, negative controls with no DCs led to no virus production, supporting that this protocol can detect specific CD4^+^ T cell responses to foreign antigens, resulting in virus production.

Taken together, these results show that recognition of self-antigens presented by DCs can induce HIV-1 expression and virus production in an MHC-II-dependent manner.

### Viruses induced ex vivo in response to autologous lysates are identical to HIV-1 variants in plasma

To test the hypothesis that the proviruses contributing to NSV are integrated into T cells reactive to specific self-associated antigens, we performed single genome sequencing of HIV-1 RNA from virus particles released following stimulation ex vivo. The sequences were then compared to viruses recovered from the plasma of participants with NSV (Figure 4).

**Figure 4.**
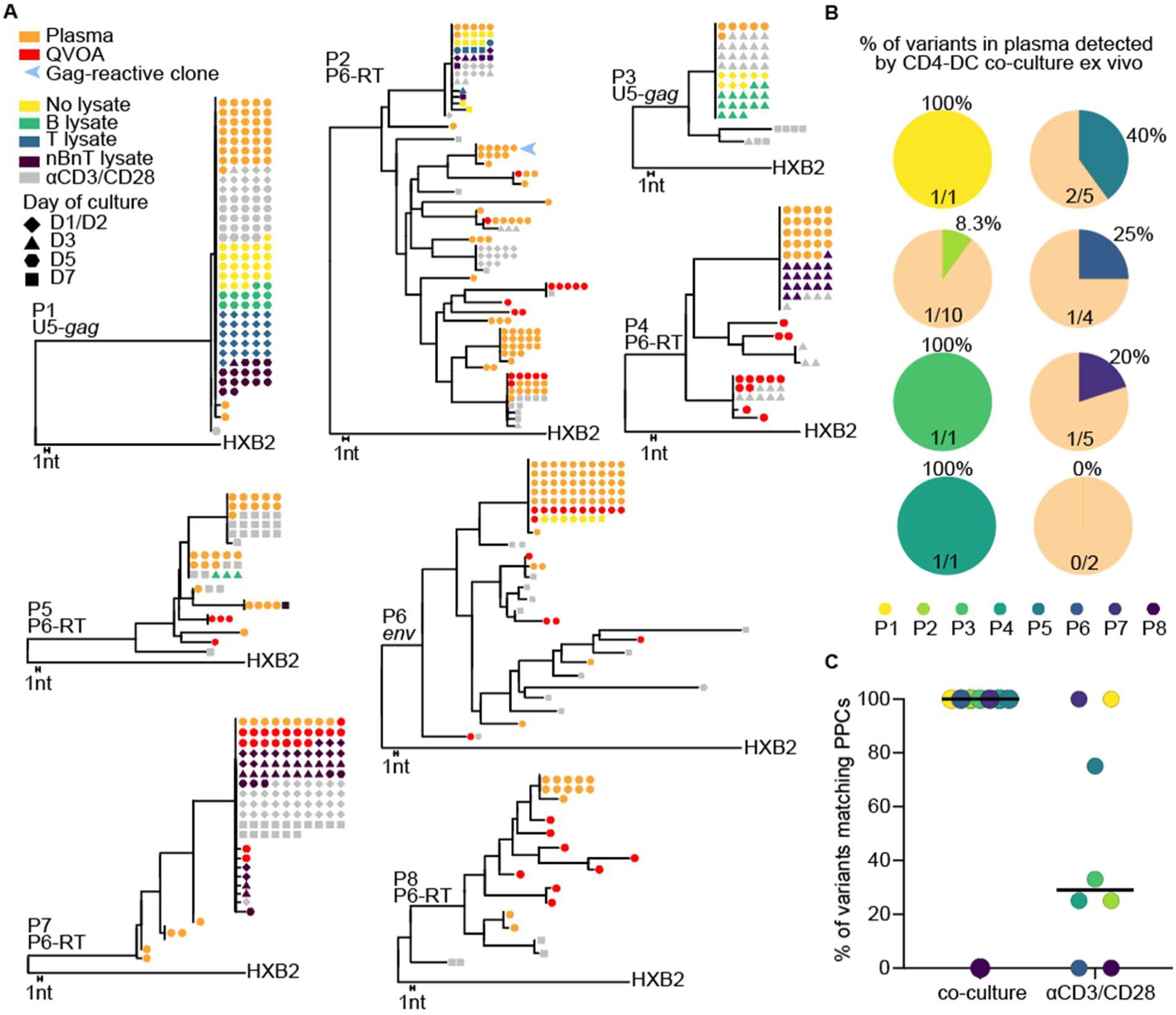
Autologous lysates induce HIV-1 variants matching virus contributing to NSV. **(A)** Maximum likelihood tree analysis of each participant with NSV. Single genome sequences of plasma viruses, viral outgrowth isolates, and HIV-1 variants recovered from the autologous assay were analyzed based on the U5-*gag* (623–1806), P6-RT (1740-3410), or *env* (7050-7980) regions. Various shapes indicate viral variants obtained from different time points of the autologous assay. Trees were rooted to HXB2. **(B)** Pie charts of the percentage of HIV-1 variants in plasma that were detected by the CD4-DC coculture. **(C)** Percentage of viral variants identical to plasma clones for each participant, based on the type of CD4^+^ T cell stimulation. Black bars indicate the median for each group.

From participant P2, we detected multiple plasma variants with variable frequencies over time, as previously described (Figure 4A) (*27*). The variant recovered from cells stimulated with autologous T and nBnT cell lysates and unpulsed DCs matched one of the predominant plasma clones. This provirus has a 21-nucleotide deletion in the MSD site and is integrated into the *DNAJB14* gene (*27*). Following ɑCD3/CD28 stimulation, 7 other viral variants were detected, 2 of which were also found in plasma (Figure 4A). In participant P4, NSV was caused by a single expanded T cell clone with a provirus integrated into the *CCND3* gene, featuring a 22-nucleotide deletion in 5’-Leader(*27*). Sequencing of the p6-RT region of viral RNA produced upon stimulation with DCs pulsed with nBnT cell lysate matched the variant causing NSV (Figure 4A).

Overall, viral variants induced by autologous lysates matched plasma viruses in 7 out of 8 participants with NSV (Figure 4). Variants produced in response to autologous lysates matched a median value of 32.5% of plasma viruses contributing to NSV. Participants P1, P3, and P4 each had a single dominant provirus contributing to viremia, and variants induced by autologous lysates resulted in a 100% match (Figure 4B and 4C). As expected, stimulation with ɑCD3/CD28 induced virus production from a broader pool of infected cells, resulting in more variants detected (Figure 4C and Figure 6C). In four individuals, the autologous lysate induced HIV-1 variants matched between 8% and 40% of viral variants found in plasma (Figure 4B). Notably, viruses induced by autologous lysates that matched plasma variants in P6 and P7 were identical to replication-competent viruses isolated from QVOA (Figure 4A). Only in participant P8, stimulation with autologous lysates did not produce virus. However, virus induced by ɑCD3/CD28 and viral outgrowth isolates also did not match PPCs, suggesting that the cells causing viremia were not circulating in blood or were not induced to express virus ex vivo.

Previous work has shown that infected clones can persist, wax, and wane over time, and this fluctuation is also reflected in the HIV-1 variants contributing to residual viremia (*30, 52, 64*). For participant P1, we had enough PBMCs to repeat the autologous stimulation ex vivo with cells from two time points (Supplementary Figure S5A and S5B). The results from the two time points were in agreement, showing production of the ADK.d22 virus upon CD4^+^ T cell culture with DCs. While enrolled in our study, P1 experienced a drop in plasma HIV-1 RNA levels to below 20 copies/mL after six years of continuous NSV (*27*) (Supplementary Figure S5C). The participant reported no change in ART or other medication; clinical records were unremarkable. Surprised by this finding, we quantified the frequency of the ADK.d22 provirus, a single PPC contributing to NSV in this participant, by integration site-specific dPCR before and after the resolution of NSV. The provirus decreased from ∼50 copies/10^6^ CD4^+^ T cells to below the limit of quantification of our assay (<1 copy/10^6^ cells) in all samples collected after viremia became undetectable (Supplementary Figure S5D). Given previous studies showing that the ADK.d22 clone represented a marked fraction of all infected cells (*27*), total LTR copies also decreased after the resolution of NSV. Moreover, when we repeated the autologous coculture experiments described above, we observed no virus release in response to cell stimulation with autologous cell lysates or even with ɑCD3/CD28 (Supplementary Figure SE). These findings suggest that the ADK.d22 clone, the primary source of NSV in this participant, had contracted to below our detection capacity in peripheral blood. This finding agrees with other longitudinal studies that have reported spontaneous resolution of detectable viremia in about a third of individuals with NSV (*65*). The mechanisms underlying such a sharp decrease in less than two months remain elusive. The functional impairment of this clonotype due to chronic stimulation –with loss of proliferative capacity– or the development of new immune pressure against the cells of this clonotype carrying the ADK.d22 provirus may explain the resolution of NSV and should be investigated in future studies.

### Proviruses induced by autologous lysates are in conventional CD4^+^ T cells

TCR avidity for self-peptides presented by MHC molecule is highly variable and determines not only thymic selection but also the fate of cells maintaining peripheral tolerance (*49, 66*). Given that the regulatory T cells (Treg) have TCRs with higher affinity to self-peptides (*67*), we investigated whether the infected clones contributing to NSV and inducible by autologous stimulation ex vivo were found in Treg or conventional (Tcon) CD4^+^ T cells. We analyzed CD4^+^ T cells based on the expression of CD25 (IL-2Rɑ) and CD127 (IL-7Rɑ) by flow cytometry and sorted CD25^hi^CD127^low^ cells (Figure 5A), which are highly enriched for *bona fide* FOXP3^+^ Tregs as previously reported (*68*). The average frequency of Treg among all CD4^+^ T cells was 5.8% (range 3.0-7.6, Figure 5B), within the range observed in people of comparable age to our participants (*69*). Digital PCR was used to quantify LTR copies (R-U5) and proviruses previously shown to contribute to NSV in P1, P2, P3, and P4 by integration-site specific assays, as previously described (Figure 5C) (*27, 70, 71*). The frequency of infected cells was comparable between Tcon and Treg cells (3238 *vs* 2313 LTR copies/10^6^ cells, p=0.68). However, the four proviruses of interest were exclusively found in Tcon cells (range 19-72 copies/10^6^ cells) and were below the detection limit in Tregs (Figure 5D). These results support the hypothesis that proviruses with spontaneous HIV-1 gene expression that maintain NSV are found in conventional T cells. This observation aligns with previous studies showing profound phenotypic heterogeneity among all infected cells (*63, 72, 73*).

**Figure 5.**
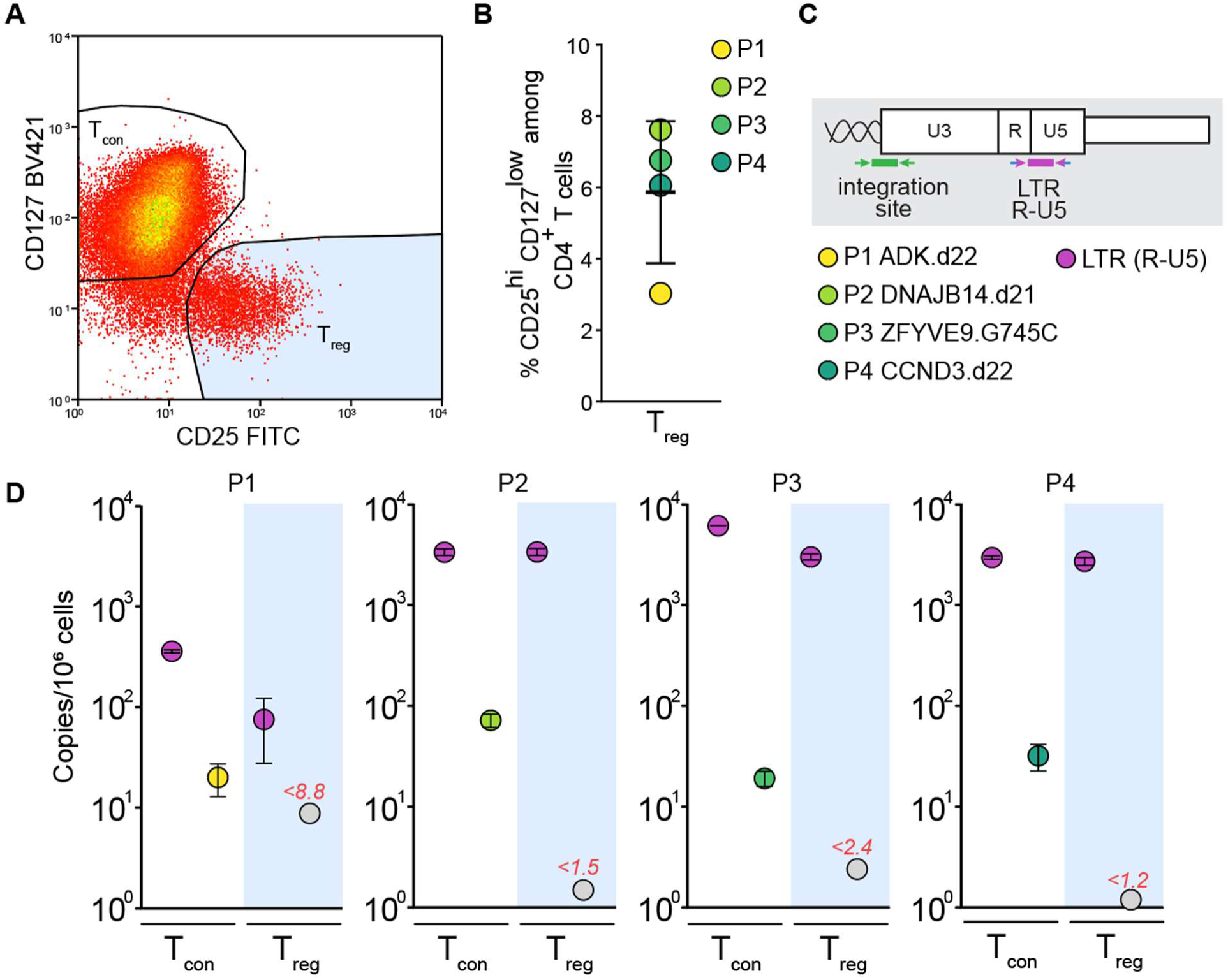
Proviruses causing NSV are not compartmentalized in regulatory T cells. **(A)** Sorting strategy for Tcon (CD127^hi^CD25^low^) and Treg (CD127^low^CD25^hi^). **(B)** Percentage of Tregs among CD4^+^ T cells in participants P1-P4. **(C)** Total LTR copies are quantified using a probe targeting the R-U5 junction. Proviruses of interest for P1-P4 are quantified using integration site-specific probes. **(D)** Frequencies of specific proviruses, expressed as copies/10^6^ CD4^+^ Tcon and Treg cells. Gray circles indicate values below the limit of detection (in red). Error bars indicate the standard error of the mean.

### Recognition of self-antigens can induce replication-competent clones in some ART-suppressed PLWH

People on ART experiencing NSV represent an extreme case of residual viremia, which can be detected by ultrasensitive assays in more than half of PLWH (*16, 17*). To investigate if virus can be released in response to autologous stimulation in PLWH with an undetectable viral load (Figure 6), we studied 5 participants with plasma HIV-1 RNA below 20 copies/mL for more than six months. Their clinical characteristics are summarized in Supplementary Table 1. Stimulation with ɑCD3/CD28 resulted in virus production from CD4^+^ T cells in all 5 participants, indicating successful detection of virions in the supernatant with nonspecific activation. In 3 out of these 5 participants (P9, P10, P13), CD4^+^ T cells produced virus in response to DC pulsed with autologous lysates (Figure 6A). In contrast, stimulation with autologous lysates from P11 and P12 did not result in virus production.

**Figure 6.**
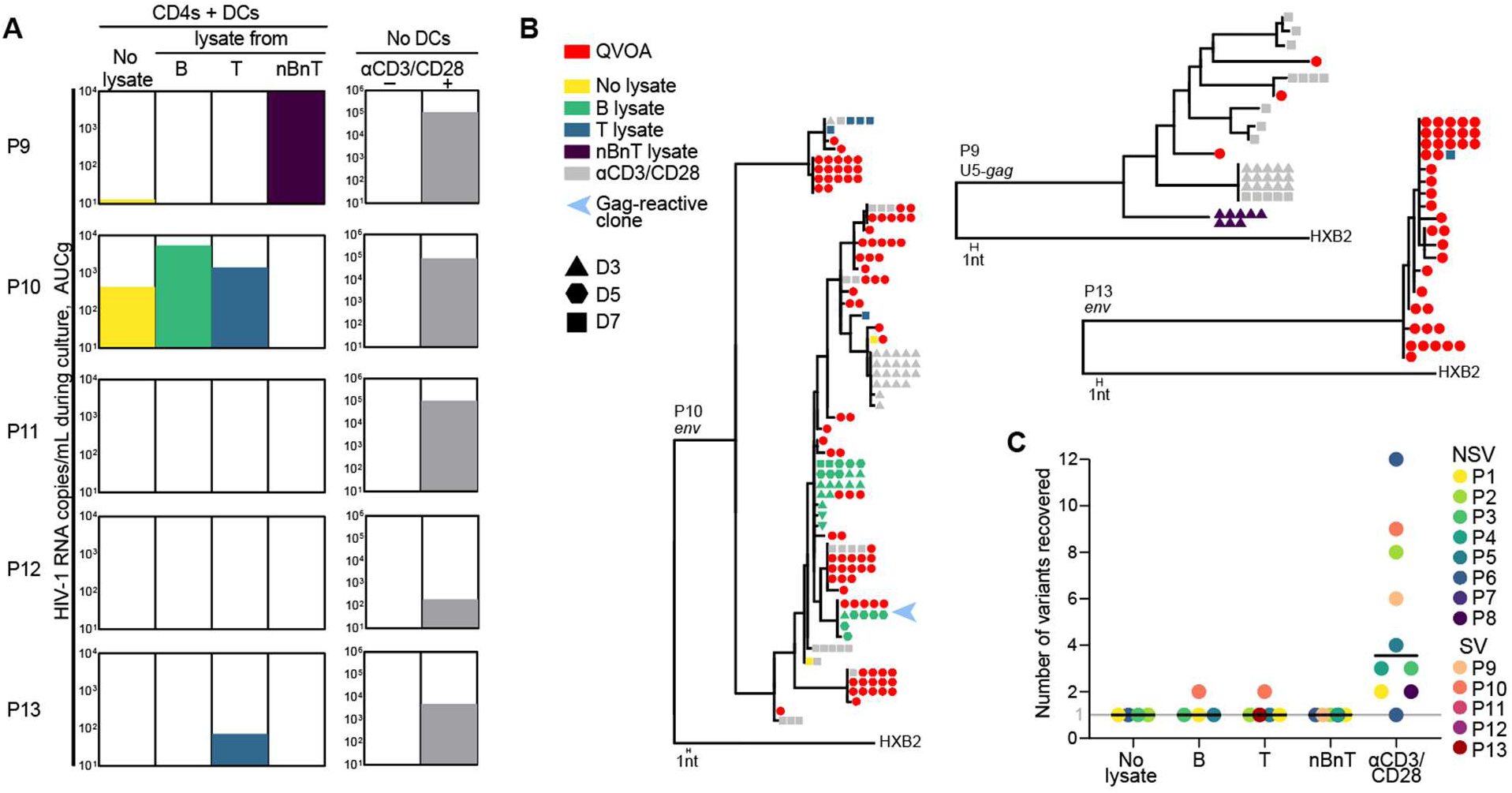
HIV-infected cells reactive to autologous lysates from replication-competent provirus. **(A)** AUCg HIV-1 RNA copies from 7 days of culture for five participants with suppressed viremia (SV). **(B)** Maximum likelihood trees for SV participants P9, P10, and P13, rooted to HXB2: U5-gag (623-1806) or env (7050-7980) regions were used for the sequence analysis. Symbol shapes indicate viral variants obtained from different culture time points. For P10, the HIV-1 variant matching a Gag-responding clone is shown with the blue arrow (see also Supplementary Figure S4). No sequences were recovered for αCD3/CD28 stimulation in P13 **(C)** Number of HIV-1 variants obtained from each group of autologous lysates (DCs only, B, T, nBnT lysates) or CD4^+^ T cells with ɑCD3/CD28. Black bars represent the mean value from each group, while the color of the dots represents the study participants (P1-P8, NSV, and P9-P13, SV). Participants without a response against autologous lysate or match to viral variants were excluded from this analysis.

In P9, limiting-dilution U5*-Gag* sequence analysis revealed that CD4^+^ T cells stimulated with nBnT lysate pulsed DCs led to a single variant that did not match any QVOA isolate (Figure 6B). In P10, B cell lysate induced two variants that matched replication-competent outgrowth virus (Figure 6B), one of which was also induced upon stimulation with Gag (Supplementary Figure S4). Two different variants were induced by unpulsed DCs, with one matching a replication-competent provirus. Due to the low intra-host diversity of viral populations from P13, it was challenging to determine whether the variant induced by DCs-pulsed with T cell lysate matched one replication-competent QVOA isolate because they belonged to the same provirus or due to overestimation of clonality (*74*).

When analyzing viral variants found in the supernatant across all 13 participants, stimulation with autologous lysate induced a median of 1 variant, reflecting the activation of rare, specific cells. Conversely, nonspecific activation with ɑCD3/CD28 resulted in a median of 5 variants (range 1-12) (Figure 6C), supporting that provirus induction and release of viral particles originated from cells that responded to specific antigens. Together, these results suggest that the recognition of self-associated antigens driving spontaneous reservoir activity is not unique to NSV participants but can also occur in some individuals on ART with undetectable viral loads. Moreover, as observed in some of the participants with NSV, clones that recognize self-associated antigens can carry inducible, replication-competent proviruses.

### Virus production induced by autologous lysates and the autoantibody reactome are similar in participants with NSV and undetectable viremia

Although virus production upon stimulation with autologous lysates varied across participants, we detected the release of virus particles in response to unpulsed DCs and DCs pulsed with B, T, and nBnT cell lysates in 10 out of 13 participants (Figure 7). When comparing the amount of virus detected in culture over time (HIV-1 RNA AUCg) between participants with NSV and those with suppressed viremia (SV), we observed no significant differences between the two groups (Figure 7A and 7B). CD4^+^ T cells left untreated or stimulated with ɑCD3/CD38 yielded similar levels of virus in cultures. Despite the limited sample size, we observed a significantly higher virus production trend in participants with NSV (p=0.054), driven mainly by stimulation with unpulsed DCs and nBnT lysate (Figure 7B). Since virus production followed a dichotomous, all-or-nothing pattern, we analyzed the proportion of participants we recovered virus from. Of the 13 study participants, 7 showed virus production from CD4^+^ T cells cultured with unpulsed DCs (5 NSV and 2 SV) (Figure 7B). In comparison, we observed virus release in response to autologous cell lysates in 7 of 8 NSV participants (87.5%) and 3 of 5 SV participants (60%). Notably, the proportion of participants with virus production with nBnT cell lysates was significantly higher in individuals with NSV (p=0.032) (Figure 7C).

**Figure 7.**
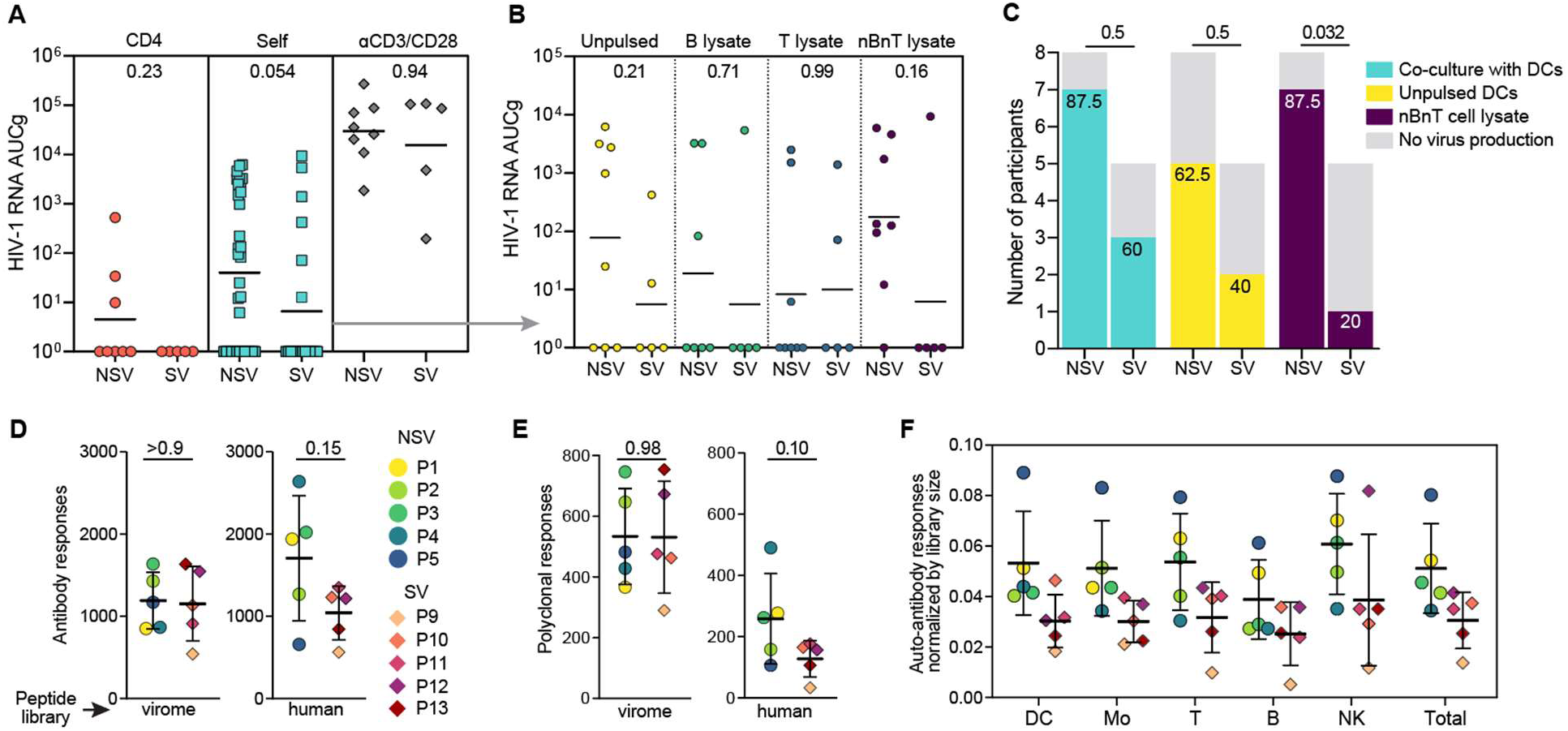
Virus production in response to autologous lysates and autoantibody reactome are similar between participants with and without NSV. **(A)** Area under the curve (AUCg) of HIV-1 detected over time from CD4^+^ T cells alone, cultured with autologous DCs, or treated with ɑCD3/CD28 between NSV and SV participants. Each dot represents AUCg value from an individual condition. We assigned a value of 1 to conditions without HIV-1 detection. The horizontal lines represent the geometric mean. **(B)** Breakdown of results from coculture experiments with CD4^+^ T cells and autologous DCs pulsed with different cell lysates. **(C)** Summary bar graph indicating the number and percentage of study participants with virus production; differences between the two groups were analyzed by Fisher’s exact test. **(D-E)** PhIP-seq analysis of antibodies binding to the virome or the human peptide libraries (see methods); each symbol indicates one study participant, and error bars represent mean and standard deviation. Differences between the two groups were tested by Mann-Whitney t-test. D shows total counts of reactive peptides, while E shows proteins against which two or more non overlapping peptides were targeted by antibodies. **(F)** PhIP-seq sub-analysis of proteins expressed by immune blood cells (Dendritic cells, monocytes, T cells, B cells, and natural killer lymphocytes) used for autologous lysates. Counts of total antibody responses were normalized by peptide library size for each cell type. Error bars indicate mean and standard deviation.

Recent advances have revealed that self-reactive antibodies (autoantibodies) play an underexplored role in immunomodulation, such as IFN-I autoantibodies mediating the risk of death from COVID-19 while reducing the severity of Systemic Lupus Erythematosus (*75, 76*). Given the role of CD4^+^ T cells in the selection and maturation of autoantibodies, these can be used to interrogate self-reactive CD4^+^ T cell responses (*50*). To investigate whether the presence of NSV is associated with a specific "antibody reactome" signature, we performed phage immunoprecipitation sequencing (PhIP-seq) (*77–80*). We tested the plasma of 10 study participants (5 NSV and 5 SV) for binding to phage-displayed antigen libraries covering the entire human peptidome and the human virome (Figure 7D and 7E; see methods for details). To validate this approach, we first assessed the antibodies against HIV-1 and four common viruses (rhinovirus, herpes simplex 1 and 2, respiratory syncytial virus, and cytomegalovirus). We used Mpox as a negative control since the study participants were enrolled before the 2022 Mpox outbreak. Broad antibody responses to these viral infections were detected in all participants except for Mpox and CMV for P2, who was known to be anti-CMV IgG negative based on clinical records (Supplementary Figure S6A and S6C). As previously reported, HIV-1-specific antibodies were mostly directed against Envelope and Gag (*81*) (Supplementary Figure S6B). Overall, we found no significant difference between NSV and SV participants in antibodies targeting viral or human proteins (Figure 7D). Nonetheless, autoreactivities were common and abundant in all participants, with an average of 1373 (sd ±653) unique reactive peptides and 193 (sd ±126) reactive proteins against which we detected polyclonal responses (i.e., autoantibodies binding to two or more non-overlapping peptides of the same protein) (Figure 7E). Interestingly, some participants with NSV had a markedly higher number of total and polyclonal autoantibody responses. This pattern was also observed after we narrowed the analysis to autoantibodies against proteins expressed by immune cells, including those we used to generate autologous lysates (Figure 7F and Supplementary Figure S7).

Taken together, these results suggest that HIV-1 proviruses can persist in cells reactive to self-associated antigens and that this phenomenon can occur in people on ART even without overt NSV. In addition, the autoantibody reactome analysis confirmed that self-reactive humoral responses are common in PLWH on ART regardless of NSV. This finding should prompt more extensive studies with matched HIV-1 negative controls, which would allow investigation of whether HIV-1 infection is associated with a specific autoantibody profile.

## DISCUSSION

Residual and nonsuppressible viremia are reflections of the same phenomenon: in vivo activation in a small fraction of the cells comprising HIV-1 latent reservoir, resulting in transcriptional activity. The clonal and non-evolving nature of this persistent HIV-1 RNA in plasma indicates virus production from latently infected cells that have proliferated and persist. The virions comprising RV and NSV cannot infect new cells because of the inhibitory effects of ART but can be detected in plasma by sensitive assays (*21–23, 26, 27, 82*). Antigen-driven cell proliferation has been recognized as a major driving force in the persistence of the latent reservoir (*12, 13, 63*). However, while the typical response for episodic exposure to foreign antigens (clonal expansion ◊ antigen clearance ◊ clonal contraction) is reflected in the waxing and waning of some infected clones, including those contributing to residual viremia (*30*), it fails to explain years of continuous viral production from specific latently infected cells (*26–28*). Thus, we hypothesized that responses to chronic viral infections, commensal microbiota, or self-antigens could drive persistent viremia. Here, we investigated the reactivity of infected CD4^+^ T cells to potential self-antigens in autologous cell lysates processed and presented by autologous DCs ex vivo. This method allowed us to exploit virus production to amplify and read out the signal from TCR:pMHC interactions of rare HIV-1-infected, antigen-responding cells. Moreover, we leveraged intra-host HIV-1 sequence diversity as a natural barcode to infer the activation of antigen-specific cells. Recent studies from our group and others have shown that physiological T cell activation with cognate antigen can induce viral gene expression (*27, 39–41*). To the best of our knowledge, this is the first study to demonstrate this process of self-antigen recognition among reservoir cells and their contribution to persistent viremia in people on ART.

Previous studies have demonstrated a weak correlation between the reservoir size and residual viremia (*83–85*). One study has shown a proportional relationship between the frequency of infected cells, HIV transcription, and virus release (*86*). Here, we found no correlation between reservoir size and NSV, as both IPDA and IUPM values fell within the range of values observed in people with undetectable HIV-1 RNA (Figure 1). Correlating persistent viremia with reservoir size is further complicated by the fact that in about a third of people with NSV, virus in plasma is derived from one or more proviruses with 5’-Leader defects that would not be measured by IPDA or QVOA (*27, 28*). In addition, we recently demonstrated that the size of infected clones contributing to viremia in P1, P2, P3, and P4 did not correlate with the amount of virus in plasma (*27*). These results, along with the previous observation that only a small fraction of cells within a clone is positive for unspliced RNA at any given time (*87*), suggest that the frequency of proviruses is not enough to explain NSV (*88*); thus, the quantification of reservoir size may not help the diagnostic work-up and clinical management of people with persistent viremia.

The site of HIV-1 integration can affect HIV-1 persistence. Several studies suggest that the pHIV-1 integration into few host genes, such as BACH2 and STAT5B, can cause insertional gene activation and drive infected cell proliferation and survival (*9, 89–92*). In addition, intact proviruses found in heterochromatic chromosomal locations may display lower inducibility, allowing them to evade immune recognition (*40, 93–95*). In our study, the proviruses with known integration sites were all found within introns of genes with variable expression in CD4^+^ T cells (*27*). This observation aligns with the study from Mohammadi et al., which found a correlation between the proximity of virus-producing proviruses to activating epigenetic marks (e.g., H3K36me3 and H3K9me3) and their HIV-1 RNA copies in plasma. Thus, integration site features may explain the differences in the levels of persistent viremia between PLWH and whether they fall within the levels of NSV (*96*).

Using DCs pulsed with autologous lysates we induced the predominant plasma clones ex vivo from the CD4^+^ T cells of 7 of 8 participants with NSV. Previous studies showed that extensive sampling is required to find matching sequences between virus in plasma and proviruses in cells due to the abundance of highly defective proviruses and the low frequency of infected clones contributing to persistent viremia (*23, 26, 27*). However, despite their rarity, we detected the variants responsible for NSV in the coculture supernatant, suggesting that the stimulation with specific self-associated antigens plays a critical role in selectively inducing the infected clones driving NSV. In most instances, virus production ex vivo was decreased or blocked by adding an ɑMHC-II antibody, suggesting that engagement of TCR:pMHC-II was necessary for latency reversal. Viruses released from CD4^+^ T cells stimulated with autologous lysates were comprised of only one or two variants and were identical to plasma viruses that sustained persistent viremia in 7 participants. While the viral outgrowth assay has been used to isolate proviruses causing NSV (*10, 26, 31*), it fails to detect non-infectious NSV variants. Global T cell activation can induce a polyclonal population of proviruses that are not spontaneously expressed in vivo and released in plasma. Therefore, our approach more specifically induced variants responsible for persistent viremia compared to these methods.

Reactivation of latently infected CD4^+^ T cells by autologous lysates was also observed in 3 out of 5 participants with undetectable viremia, with 2 cases matching viral outgrowth isolates. This finding suggests that proviruses from latently infected CD4^+^ T cells can be reactivated by self-associated antigens regardless of the level of persistent viremia, reinforcing the idea that NSV is an extreme example of the same phenomena that drive RV and likely occur in most people on ART (*29*). Furthermore, in participants P2 and P10, virus production was induced by both autologous lysates and HIV-1 Gag. In P2, the proviruses induced by HIV-1 Gag also matched the virus found in plasma. This finding supports the idea that antigens from chronic viral infections – such as HIV-1 itself – can contribute to spontaneous reservoir activation (*41*) and persistent viremia, leading to the hypothesis that in some individuals, the release of viral particles from Gag-reactive CD4^+^ T cells could result in positive feedback that sustains persistent viremia over time.

The presence of HIV-1-infected CD4^+^ T cells reactive to self-associated antigens should not be interpreted as a reflection of pathogenic clones driving autoimmune disease. Seminal studies have shown that T cells exhibiting overt reactivity to self-ligands are negatively selected in the thymus and further removed from the conventional T-cell repertoire by deletion, anergy, or differentiation into alternative lineages. However, these processes are imperfect, and peripheral self-reactive cells are common (*66, 97, 98*). Su *et al.* identified memory CD4^+^ T cells specific for self-antigens in healthy adults, at frequencies ranging between 1 to 10 cells per million CD4^+^ T cells, similar to those of virus-specific cells in unexposed individuals (*47*). This frequency of cells recognizing self-antigens is likely the result of a compromise between physiological selection and inherent immunity to foreign pathogens (*46*). The number of foreign pMHC that T cells must recognize (>10^15^) to provide effective immunity is much greater than the number of T cells available (10^12^) (*46*). Therefore, to provide comprehensive coverage, the immune system must rely on cross-reactivity and extensive TCR promiscuity, which is even more pronounced for CD4^+^ T cells due to the open ends of the MHC-II cleft.

Our study has important implications for understanding nonsuppressible viremia and reservoir persistence on ART. We demonstrate that self-associated antigens play a role in the activation and likely the expansion of infected clones, as previously demonstrated for microbial antigens (*12, 13, 41, 52, 63*). The availability of ubiquitous self-associated antigens could result in frequent encounters of TCR:pMHC and drive T cell activation and persistent virus production despite effective ART. In 4 participants (P6, P7, P10, P13), we matched viruses from cells reactive to autologous cell lysates to replication-competent viruses. Rebound-competent viruses are typically clonal and can be found as transcriptionally active prior to ART interruption (*33, 99*). Although rare, infected cells that are reactive, or cross-reactive, to self-associated antigens could carry proviruses and first rekindle infection upon ART cessation, leading to viral rebound. While self-peptides are constitutively presented by MHC-II molecules of professional APCs (*100*), self-reactive cells are rare. Given that the frequency of infected cells that persist on ART and can release viral particles is approximately 1-10 per million resting CD4^+^ T cells (*5, 7, 54, 101*), self-reactive cells carrying proviruses that could cause NSV even less frequent. This rarity may explain why NSV is relatively uncommon even though its underlying mechanism – virus expression from clonally expanded infected cells – operates in all PLWH on ART. Moreover, additional factors play a role in the development of NSV, such as the size of the infected clone, proviral location, and host immune pressure (*26–28*). Despite the limited sample size, we showed that these HIV-1-infected cells reactive to autologous antigens are part of pervasive immune responses to self-antigens, detectable even in individuals without overt autoimmune diseases, as our autoantibody reactome analyses supported. Future studies on larger cohorts and matched controls should investigate whether the presence of specific autoantibodies (e.g, interferon pathways, homeostatic cytokines) can affect HIV-1 progression and reservoir persistence. Additionally, although self-antigen recognition may drive virus expression and chronic inflammation, the reverse relationship is also possible. Hyper-inflammatory conditions can lower the threshold for autoimmunity (*102*) and lead to more frequent HIV-1 reactivation. Given that persistent inflammation affects PLWH even after long-term ART, future studies should investigate whether resolving NSV would decrease inflammation or if NSV is merely a consequence of self-reactivity and immune activation. The present study has some limitations. Several possibilities could explain the lack of detectable virus production from infected CD4^+^ cells in some study participants. These include a low frequency of infected self-reactive cells, the antigen of interest not being present in the lysate, or being present at very low concentrations. Since we only sorted PBMCs based on lineage markers, tissue-specific cells and their antigens were not tested. Additionally, the lysates used for the assay likely contained only soluble cellular proteins, excluding insoluble proteins such as membrane-associated or hydrophobic proteins (*103, 104*). Most importantly, we did not identify the self-associated antigen at the protein or peptide level, which would provide definitive evidence of CD4^+^ T cell self-reactivity (*105, 106*). Due to sample limitations, we were unable to pursue i) antigen discovery experiments that require more than 10^8^ cells for MHC-II ligandome analyses (*107*), and ii) single-cell approaches that would allow recovering the TCR alpha and beta chains from infected clones causing NSV (*63, 99*), which can be used to screen their reactivity against large pMHC-II libraries (*108*). Although our data suggest that the clones driving NSV are conventional rather than regulatory T cells, knowing the specific pMHC ligand would allow the phenotypic characterizations of these infected cells.

In conclusion, we demonstrated that HIV-1-infected CD4^+^ T cells can be activated by DCs pulsed with autologous cell lysates, resulting in latency reversal and production of viral particles from the same proviruses that cause NSV. Our work provides new insights into the spontaneous transcriptional activity of the HIV-1 reservoir, which sustains persistent viremia, may contribute to deleterious immune activation, and, in some individuals, to viral rebound upon ART interruption. Finally, these findings prompt future research efforts to better characterize the reactivity of infected clones responding to chronic antigens, as it would contribute to the development of individualized approaches targeting transcriptionally active proviruses and improve clinical outcomes for PLWH.

## MATERIALS AND METHODS

### Study Participants

Characteristics of all study participants are provided in Supplementary Table 1. Participants with NSV were referred from The John G. Bartlett Specialty Clinic, Johns Hopkins University and Clinique I.D. of Saint-Jérôme (Quebec, Canada). Criteria for NSV study participants include (a) initial ART-suppression of viremia with VL <20 copies/mL by clinical assays followed by (b) sustained or intermittent VL of >20 copies/mL for at least 12 months, and (c) lack of issues in adherence assessed by the clinical care providers and drug resistance to the current regimen. Changes or intensification of ART regimen did not exclude participants from the study. Peripheral blood samples (up to 180mL) were collected from at least two time points. Plasma and peripheral blood mononuclear cells (PBMCs) were isolated as described below.

ART-suppressed PLWH samples were obtained from University of Pennsylvania. Selection criteria for ART suppressed PLWH include (a) viral suppression with VL <20 copies/mL for more than 6 months and (b) CD4 count >400 cells/uL. Participants from University of Pennsylvania underwent leukapheresis at a single time point. PBMCs were isolated and viably frozen.

### Sample Processing

Whole blood samples obtained from study participants were spun at 400xg for 10min at room temperature and the plasma fraction was spun again at 1200xg for 10min at room temperature before freezing at -80C. Blood samples were processed and PBMCs were isolated using density centrifugation on a Ficoll gradient as previously described(*27*).

### Generation of autologous lysates

Viably frozen PBMCs were thawed in RPMI 1640 medium with 50% Fetal bovine serum (FBS) and spun down to remove DMSO, followed by rest for 4 hours in RPMI with 10% FBS. Cell pellets were resuspended with 75uL of wash media and incubated with 10uL of TruStain FcX (BioLegend) for 10 min, at room temperature. Cells were then stained with 5uL of CD19 BV421 (Clone HIB19, BioLegend), 5uL of CD20 BV421 (Clone 2H7, BioLegend), and 10uL of CD3 APC (Clone UCHT1, BioLegend) followed by incubation on ice for 30min. After incubation, cells undergo two rounds of washes with wash media. Dead cells were excluded by propidium iodide and cells stained with single fluorophore-labeled antibodies were used to set sort gates. A representative sort gating strategy and sort logic is provided in Figure 2 and Supplementary Figure S2. Cells were sorted using MoFlo Legacy or the XDP cell sorters. Sorted cells fractions were pelleted and resuspended with 1X PBS at 50 million cells/mL. Cell suspensions were then lysed by undergoing 6 freeze-thaw cycles on isopropanol and dry ice followed by incubation in a water bath at 37°C, respectively. Lysates were then centrifuged at 15,000xg for 10min at 4°C and the supernatant was aliquoted and frozen at -80°C. Protein concentration was quantified using Pierce Rapid Gold BCA Protein Assay Kit (ThemoFisher).

### Monocyte-derived Dendritic Cells (DCs) and coculture with CD4^+^ T Cells

Viably frozen PBMCs were thawed as previously described followed by CD14^+^ monocyte isolation (Miltenyi). CD14^+^ cells are then resuspended at 10^6^ cells/mL of X-VIVO media (Lonza) supplemented with 10% human serum (Sigma Aldrich), 400IU/mL IL-4 (R&D Systems), and 1000IU/mL GM-CSF (R&D Systems). Cells were plated in 48 well plates (5 x 10^5^/well). After two days, 5ug (10ug/mL) of lysates are added to wells containing DCs. On day five, CD4^+^ T cells were negatively isolated (Stemcell) and resuspended to 10^6^ cells/mL with TexMACS (Miltenyi) supplemented with 10% human serum, 10uM DTG (GSK1349572, Selleck Chemicals), 1X Antibiotic Antimycotic Solution (Sigma Aldrich), and 20IU/mL IL-2 (R&D Systems). After carefully removing the media from wells containing DCs, DCs were preincubated with 1ug/mL of either isotype control (Clone G155-178, BD) or MHC-II block (Clone Tü39, BD) for at least 2hrs before CD4^+^ cells were added at 1:1 cell ratio. In addition, wells were supplemented with 5ug of autologous lysates once at the start of coculture. Wells containing CD4^+^ T cells only were activated using 25uL of ɑCD3/CD28 Dynabeads per 10^6^ cells(Thermo Fisher). 200uL of supernatants were collected on days 1, 2, 3, 5, and 7, and supplemented with 200uL of fresh media. Collected supernatants are then stored at -80°C until further processing.

### Isolation of plasma and supernatant RNA

Upon thawing both plasma and supernatant, samples were briefly spun at 2,700xg for 15 min, at 4°C and transferred to a fresh Eppendorf Low-Bind tube and spun at 21,000xg, 2hr, 4°C. RNA from viral pellets was extracted as previously described(*109*).

### cDNA synthesis

Viral RNA samples were resuspended in 20uL of RNA buffer (5mM Tris HCl, 1mM DTT, 2.5mM RNaseOUT) and immediately used for reverse transcription using SuperScript III (Thermo Fisher). For cDNA synthesis of PolyA RNA measurements, 10uL of RNA were used as input with a mixture of 2.5uL of 10mM dNTP, 2.5uL 50uM oligo dT (IDT), 2.5uL 50uM random hexamers (IDT) and followed procedure described below. For gene-specific cDNA synthesis, a mixture of 20uL RNA, 2.5uL of 2uM gene specific primers and 2.5uL of 10mM dNTP were heated to 65°C for 10 min then placed on ice for 1 min. A reaction mixture of 10uL of 5x First Strand Buffer, 13.5uL of water, 0.5uL of 0.1M DTT, 0.5uL of SuperScript III RT, and 0.5uL of 40U/uL RNase Out is then added for a final volume of 50uL reaction. The mixture is then heated at 50°C for 55 minutes and inactivated at 85°C for 10 min.

### Droplet digital PCR targeting RPP30 and HIV-1 Polyadenylated RNA

Quantification of housekeeping gene, RPP30, to account for shearing was performed as previously described(*110*). Quantification of HIV-1 PolyA RNA was performed on cDNA using primers and FAM-labeled probe as previously described(*27*): forward primer GCCCTCAGATGCTRCATATAA, reverse primer TTTTTTTTTTTTTTTTTTTTTTTTTGAAG, and probe 56-FAM/TGCCTGTAC/ZEN/TGGGTCTCTCTGGTTAG/3IABkFQ (IDT). ddPCR experiments were performed using BioRad QX-200 system. HIV-1 RNA was normalized as copies/mL of supernatant. AUC calculations were made using "area under the curve with respect to ground" (AUCg) method as previously described(*62*).

### Analysis of HIV-1 sequences

The cDNA synthetized with a gene-specific primer was serially diluted with 10mM Tris HCl to reach limiting dilution and used as input for single genome sequencing as previously described(*13*). Sanger sequencing (Azenta) from positive PCR reactions were processed and raw data were analyzed using Geneious. Sequence contigs were then aligned using ClustalW(*111*). Maximum likelihood trees were constructed in MEGA 7.0 with an HKY substitution model, gamma distributed sites, 1000 bootstrap replications, and HXB2 as the outgroup reference sequence.

### Quantitative Outgrowth Assay (QVOA)

QVOA from total CD4^+^ T cells were performed as previously described. Sequencing of supernatant from p24^+^ wells were performed as described above(*112*).

### Proliferation and activation assay with ɑMHC-II

CD8-depleted PBMCs from an individual with known CMV/HIV co-infection were stimulated either with 5ug CMV lysate or 1ug/mL of PHA. Wells were then either supplemented with 1ug/mL of isotype control or ɑMHC-II block. For the activation-induced marker (AIM) assay, 1ug/mL of CD40 blocking antibody (Clone HB14, Miltenyi) was supplemented into media to prevent CD154 down-regulation. After 24 hours of activation, cells were resuspended with 75uL of wash media and stained with 2uL of CD69 FITC (Clone FN50, BioLegend), 2uL of CD154 BV421 (Clone 24-31, BioLegend), and 2uL CD137 PE (Clone 4B4-1, BioLegend) for activation markers. Samples were additionally stained with 1:1000 dilution of Fixable Viability Dye eFluor 780 (ThermoFisher), 3uL of CD3 APC (Clone UCHT1, BioLegend), and 3uL of CD4 BV605 (Clone OKT4, BioLegend). For the proliferation assay, cells were stained with CellTrace Violet (Thermo Fisher) according to manufacturer’s protocol and cells were left for 7 days without media change. Stained samples were acquired on Cytek Aurora.

### Analysis and sorting of regulatory T cells

Viably frozen PBMCs were thawed and rested for 4 hours in RPMI with 10% FBS. CD4+ T cells were isolated by negative selection (StemCell) and incubated with TruStain FcX (BioLegend) for 10 min, at room temperature. Cells were then stained as described above with 5uLof anti-CD127 BV421 (Clone A019D5, BioLegend), 5uL of anti-CD25 FITC (Clone M-A251, BioLegend), 3uL of anti-CD3 APC (Clone UCHT1, BioLegend), 4uL of anti-CD4 PE-Cy7 antibody (Clone RPA-T4, Biolegend), followed by incubation on ice for 30min. Dead cells were excluded by propidium iodide. Cells were sorted using MoFlo Legacy or the XDP cell sorters. Sorted cells fractions were pelleted and stored at -80°C until used for genomic DNA isolation as previously described (*27*). Genomic DNA was used directly as input for integration site-specific digital PCR, as previously described(*27*). Total LTR copies (R-U5 junction) and proviruses of interest (ADK.d22, DNAJB14.d21, ZFYVe9.G745C, and CCND3.d22) were normalized to copies/10^6^ cell equivalents based on RPP30.

### Analysis of HIV-1 infection frequency in monocytes versus CD4+ T cells

Viably frozen PBMCs from four participants (P1, P2, P3, P5) were thawed, rested, and stained as described above for the generation of cell lysates, but with 3uL of anti-CD3 APC antibody (Clone UCHT1, BioLegend), 5uL of anti-CD19 FITC antibody (Clone HIB19, BioLegend), 5uL of anti-CD14 BV421 antibody (Clone M5E2, BioLegend), and 4uL of anti-CD4 PE-Cy7 antibody (Clone RPA-T4, Biolegend). Dead cells were excluded using propidium iodide. Cells stained with single fluorophore-labeled antibodies were used to set sorting gates. Cells were sorted using the Beckman Coulter XDP cell sorter. A representative gating strategy and sorting logic is provided in Figure S2D. We isolated between 2.1 and 3.5 million CD14^-^/CD19^-^/CD3^+^/CD4^+^ cells and between 0.6 and 3.3 million live CD3^-^/CD19^-^/CD14^+^ cells. Sorted cell pellets underwent genomic DNA isolation genomic DNA extraction as previously described(27). The frequency of total HIV-1 DNA was measured by LTR copies, as described above, and intact proviruses were measured with the Intact Proviral DNA assay as previously described(*110*). HIV-1 copies were normalized by RPP30 as described above.

### Phage immune precipitation and sequencing (PhIP-seq)

Plasma samples were analyzed by PhIP-seq as previously described(*77, 113–115*). Briefly, we used a mid-copy T7 bacteriophage display library spanning i) the human virome, which consists of 106,678 56-mer peptides from more than 200 viruses that infect humans, and ii) the human peptidome which consists of 90-mer peptide tiles and 56-mer peptide tiles (259,345 90-mers and 14,862 56-mers for 274,207 total peptides) from 35,982 proteins; the overlap for adjacent peptide tiles is 45-aa and 28-aa for 90-mer tiles and 56-mer tiles respectively. Plasma samples were diluted 1:20 in PBS and 4uL were mixed with 1mL of phage libraries at 10^5^-fold coverage per library. After overnight incubation at 4°C in 96 deep well plates, antibody-bound phage was immunocaptured using magnetic beads coated with protein A and protein G. After bead washing, DNA tiles in the immunocaptured phage were amplified by PCR using primers containing sample-specific barcodes and P5/P7 Illumina sequencing adapters. PCR products were sequenced using a NovaSeq 6000 SP flow cell (Illumina, San Diego, CA) to determine the amino acid sequences of the antibody-bound peptides. Data from mock immunoprecipitation reactions (no plasma) were used for normalization as described previously(*116*). Two samples were analyzed in duplicates to assess sample variation and reproducibility (Supplementary Figure S7). For a peptide to be considered reactive, the read count needed to exceed 10, the fold change needed to be at least 5 and the p value of differential abundance needed to be lower than 0.001 (fold-changes and p values were calculated using the EdgeR software)(*117, 118*). Fold change values that fulfilled these three criteria were referred to as hits fold change (HFC) and the HFC of any peptide that failed at least one criterion was set to 1 (unenriched with respect to mock IP conditions). These thresholds were established based on heuristic analysis of duplicate samples, to optimize the sensitivity and reproducibility of the assay. We expect that HFC values roughly correlate with antibody abundance and/or affinity. Antibodies targeting non-overlapping peptides of the same protein were defined as "polyclonal responses".

Human peptidome hits were parsed for each participant, duplicates due to polyclonal responses were removed, and compared to publicly available dataset of gene expression enrichment in immune cell types (*119*) were obtained from The Human Protein Atlas (https://www.proteinatlas.org). A list of cell lineage enriched genes was obtained for granulocytes (1006 genes), monocytes (132 genes), dendritic cells (232 genes), NK-cells (65 genes), B-cells (301 genes), and T-cells (460 genes). Autoantibody hits that are enriched in more than one cell type are indicated on the heat map (Supplementary Figure S7).

### Study Approval

The Johns Hopkins University and the University of Pennsylvania Institutional Review Board (IRB) approved this study. All participants provided with written informed consent before enrollment into study. P4 provided written consent to IRB-approved study from laboratory of Dr. Cécile Tremblay at the Centre Hospitalier de l’Université de Montréal (CHUM), Canada.

### Statistical Analysis

Descriptive statistics, normality test, and 2-tailed student T-test were used to determine statistical significance using GraphPad Prism v8.0. A p-value of <0.05 was considered significant unless otherwise stated.

## Supporting information

Supplementary Data on PHIPseq

## Data Availability

HIV-1 sequences are available on GenBank (<u>accession numbers pending from GenBank</u>). Integration site sequencing data can be found on the NCI Retrovirus Integration Database (https://rid.ncifcrf.gov/).

## List of Supplementary Materials

-Fig. S1 to S7 for multiple supplementary figures

-Table S1

-Data file S1

## Acknowledgements

We deeply thank the study participants and their families for the commitment in volunteering in this study. We thank Joel N. Blankson for discussion leading to this work, Sylla Mohamed for assistance in sample preparation, Kenneth Lynn for clinical support, and Alicia Edwards for administrative support.

## Author contributions

F.W and F.R.S. conceptualized the study. F.W, M.M., and F.R.S. designed the experiments and performed analyses. F.W., M.M., F.D., N.L.B, A.C.C., S.B., V.H., and F.R.S. performed experiments. F.R.S, carried out integration site-specific digital PCR and related analyses. H.Z. conducted cell sorting experiments. S.J. and H.B.L. performed PhIP-seq experiments and related analyses. J.L., A.S., S.P., F.C., A.C, C.T., C.J.H., M.Z, and L.J.M. enrolled the study participants and gathered clinical data. J.D.S. and R.F.S. helped supervise the project and provided critical input for study design, data analysis, and interpretation. F.W., J.D.S., R.F.S, and F.R.S wrote the manuscript with feedback from all authors.

