## Supplementary Materials for "Proviruses in CD4^+^ T cells reactive to autologous antigens contribute to nonsuppressible HIV-1 viremia"

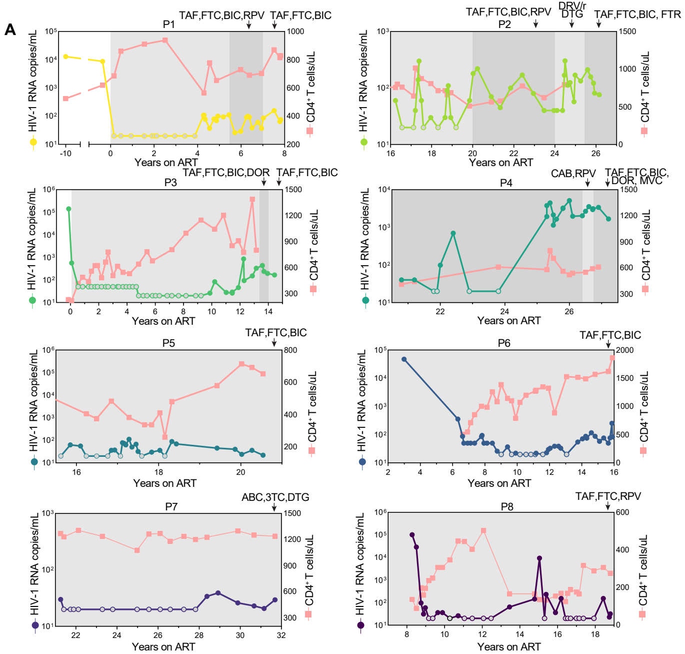


**Supplementary Figure S1. Clinical history of NSV participants.** (**A**) CD4+ T cell counts and plasma HIV-1 RNA copies over time for all NSV participants. Gray circles represent values below the limit of quantification for the assay. Light gray shaded area represents time on ART. Dark gray shaded area represents ART intensification with more than 3 drugs. BIC, bictegravir; FTC, emtricitabine; TAF, tenofovir alafenamide; DOR, doravirine; MVC, maraviroc; ABC, abacavir; DTG, dolutegravir; 3TC, lamivudine; RPV, rilpivirine; TDF, tenofovir disoproxil fumarate; ATZ/r, atazanavir boosted with ritonavir; DRV/r, darunavir boosted with ritonavir; CAB, cabotegravir; FTR, fostemsavir

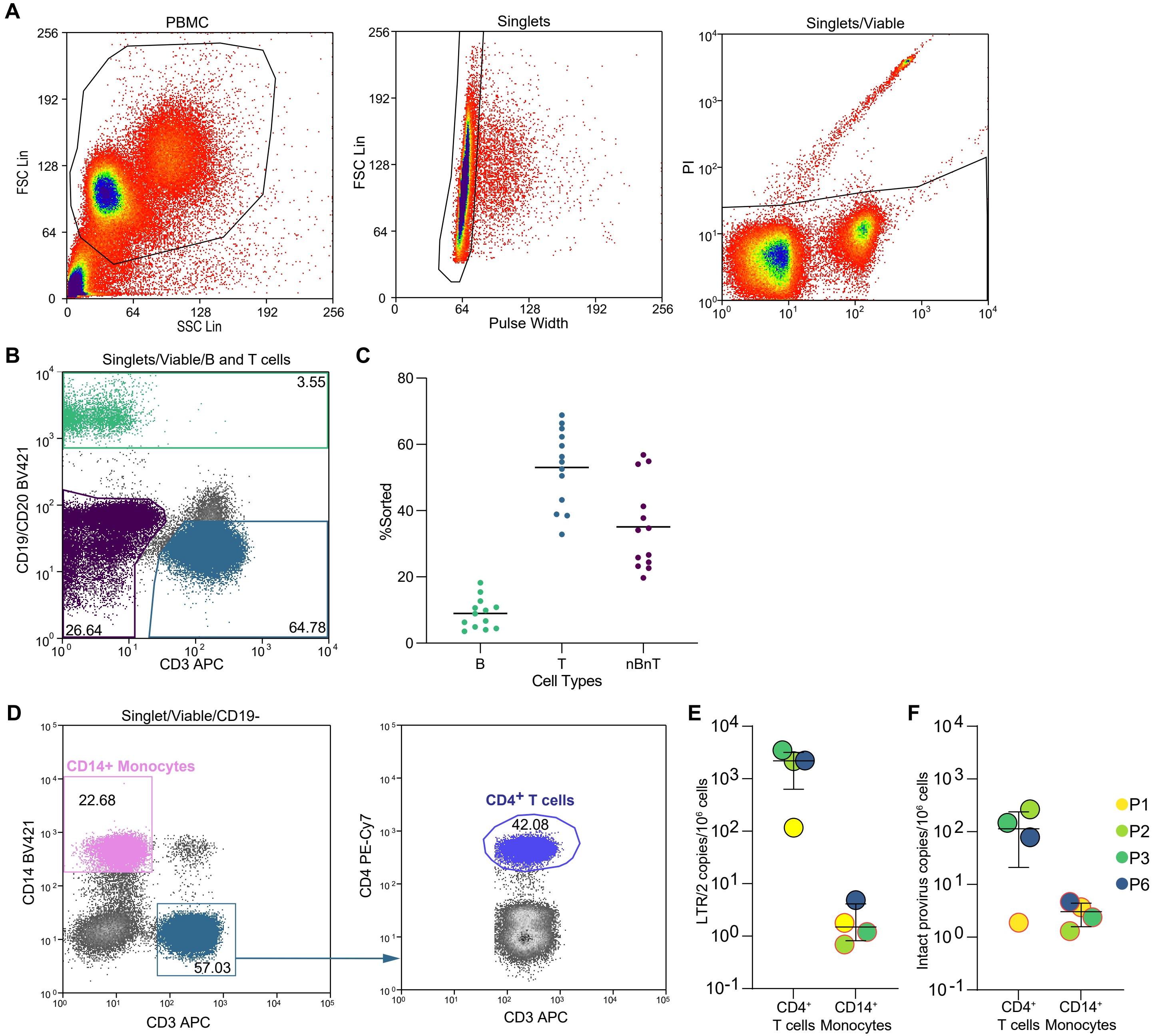


**Supplementary Figure S2. Sorting strategy of PBMC lineages for the generation of autologous lysates.** (**A**) Gating strategies to sort live PBMCs. Standard gating based on forward versus side scatter, followed by singlet gating of forward height and pulse width. Subsequently gating for single live cells using propidium iodide (PI) staining. (**B**) Sorting for B, T, and non-B-non-T (nBnT) immune cell types using CD19, CD20, and CD3 markers. Percentages of each immune cell type is shown within the selected gates. (**C**) Percent of each cell type sorted, with each dot representing a single study participant. **(D)** Gating strategy to sort P1, P2, P3, and P5 PBMCs for CD14$^+$ monocytes and CD3$^+$CD4$^+$ T cells. Cells gated for CD3$^+$ were then gated on CD4$^+$. **(E)** Sorted CD3$^+$CD4$^+$ T cells and CD3$^-$CD14$^+$ monocytes were then analyzed by digital PCR to quantify infected cell frequency by LTR (LTR copies/2) and **(F)** intact proviral DNA assay**.** Circles highlighted with a red border indicate values below the limit of detection, calculated based on the total number of RPP30 genome equivalents assayed.

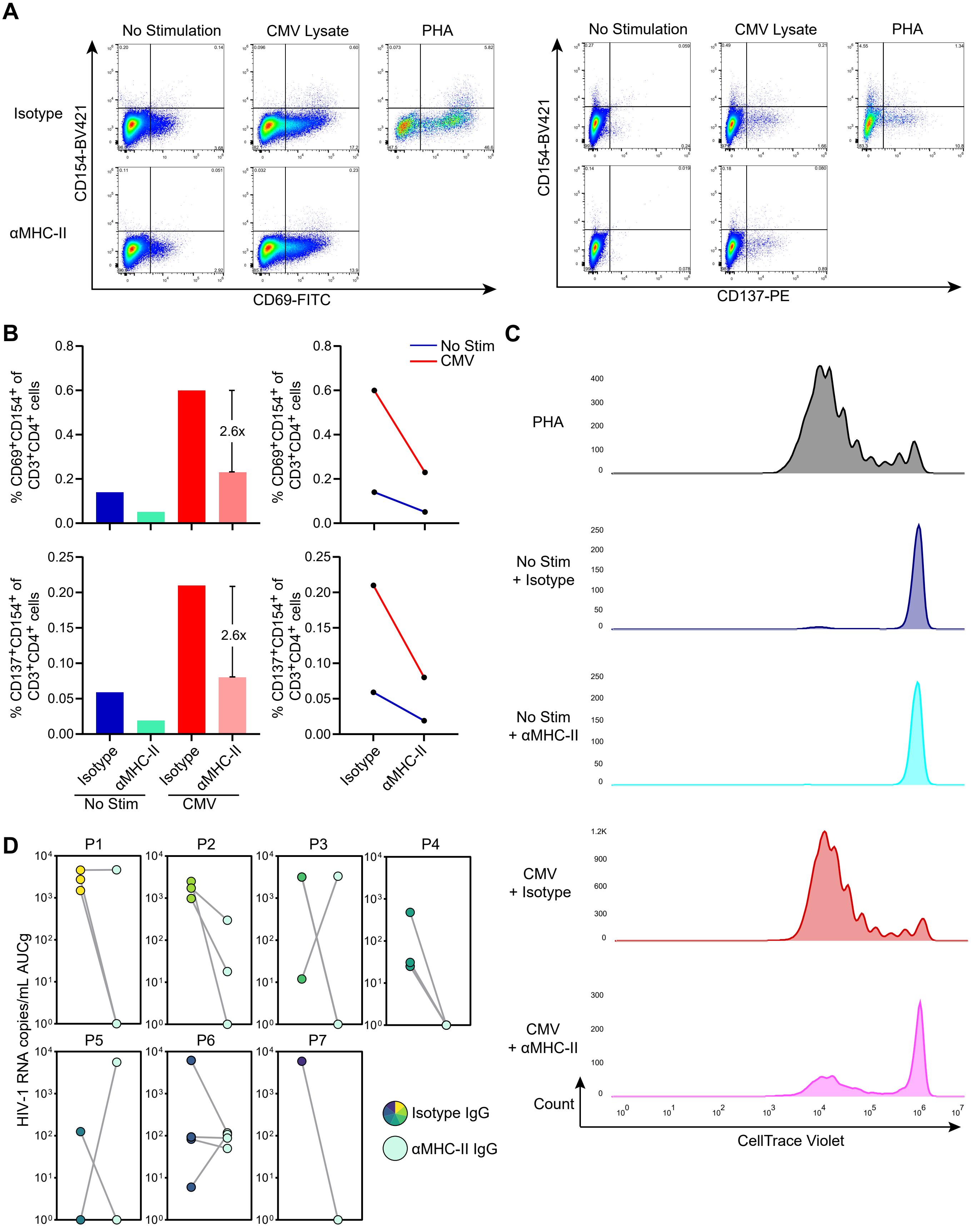


**Supplementary Figure S3. αMHC-II antibody reduces activation and proliferation of CD4+ T cells stimulated with CMV lysate.** (**A**) Flow cytometry plots showing CD154-BV421 versus CD69-FITC or CD137-PE after no stimulation, stimulation with CMV lysate, and stimulation with PHA, with either isotype control or αMHC-II block. (**B**) Percentages of CD69+CD154+ and CD137+CD154+ in CD3+CD4+ T cells measured in the stimulation assay with either isotype control or αMHC-II block. (**C**) Histogram of Cell Trace Violet intensity in cells after no stimulation, stimulation with CMV, and stimulation with PHA, with either isotype control or αMHC-II block. **(D)** Charts plotting HIV-1 RNA copies/mL AUCg for isotype control versus αMHC-II in P1, P2, P3, P4, P5, P6, and P7.

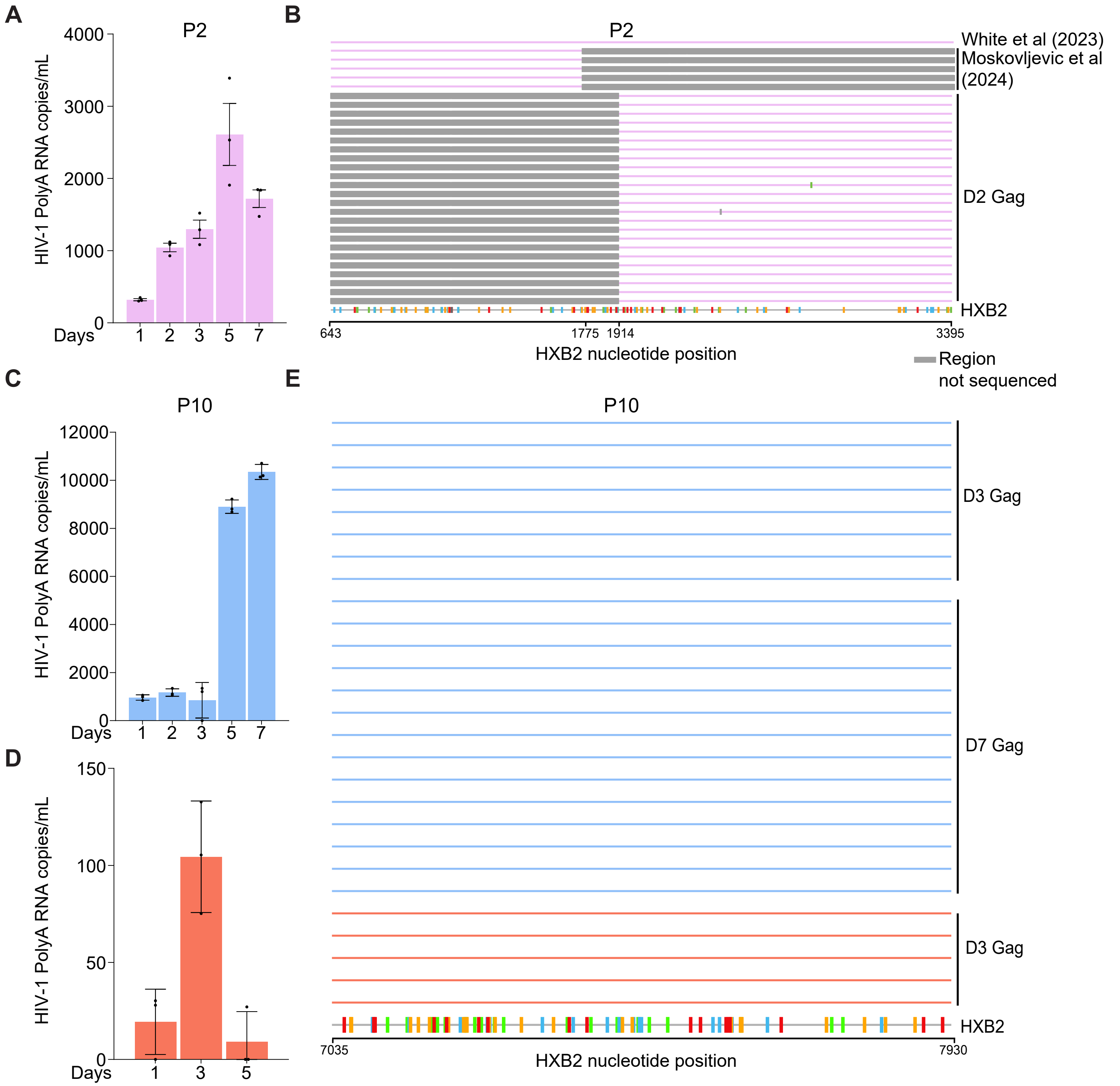


**Supplementary Figure S4. Repeated experiments of ex vivo stimulation with HIV-1 Gag protein.** (**A**) From P2, analysis of supernatant after stimulation of CD4⁺ T cells using DCs pulsed with Gag. Measurements for HIV-1 PolyA copies were obtained using ddPCR. (**B**) Comparison of the u5-RT reference sequence of a persistent plasma clone (RRM1.T745A) to the sequences recovered from the culture supernatant of CD4s stimulated with Gag from the experiment in (**A**) and from the restimulation of Gag-reactive cells expanded ex vivo from Moskovjlevic et al. (**C-D**) From P10, analysis of supernatant after stimulation of CD4⁺ T cells using DCs pulsed with Gag. Measurements for HIV-1 PolyA copies were obtained using ddPCR. (**D**) HIV-1 PolyA RNA copies for the repeat experiment of ex vivo stimulation with DCs pulsed with Gag in participant P10. (**E**) Highlighter plot of *env* sequences obtained from (**C**) and (**D**).

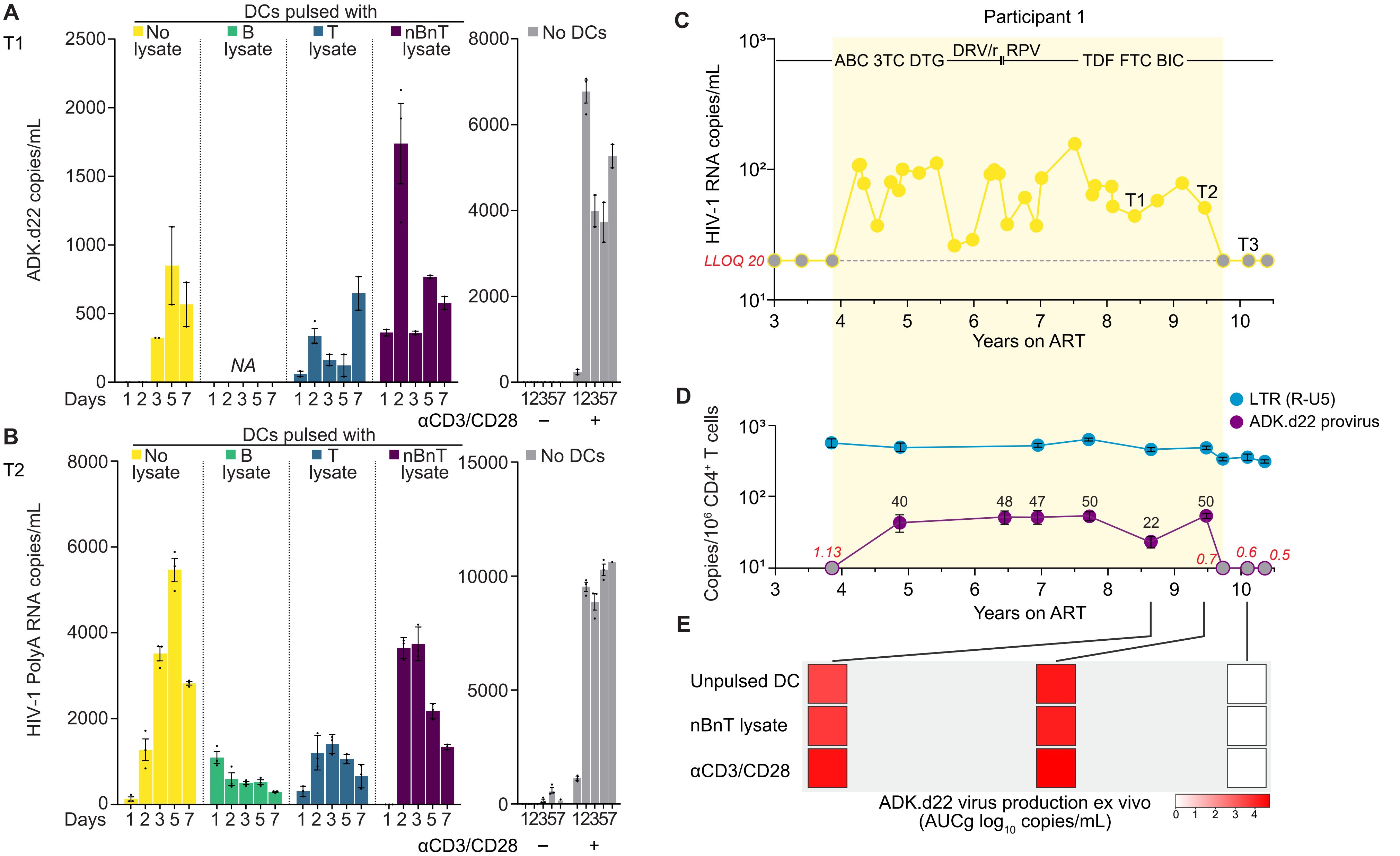


**Supplementary Figure S5. Longitudinal analyses of NSV and its underlying clonal dynamics from participant P1. (A)** Ex vivo stimulation of CD4⁺ T cells using DCs unpulsed, or pulsed with autologous lysates from P1. CD4⁺ T cells alone were also cultured alone with or without αCD3/CD28 (grey). Supernatants were analyzed using ADK.d22-specific assay harnessing the 5'Leader 22-nucleotide deletion. **(B)** Ex vivo stimulation of CD4⁺ T cells from a second time point; HIV-1 RNA was quantified by polyA RNA assay and ADK.d22 identity was confirmed by single genome sequencing (Figure 4). (**C**) Plasma HIV-1 RNA measurements show six years of continuous NSV, unaffected by ART modifications; grey circles indicate values below the lower limit of quantification (20 copies/ml, dashed line) and no HIV-1 RNA target detected. (**D**) Frequency of all LTR copies (blue) and ADK.d22 (purple) per milion CD4⁺ T cells. The limit of detection is indicated in red for time points with values equal to zero (grey circles). (**F**) Quantification of the variant contributing to NSV by stimulating CD4⁺ T cells ex vivo with unpulsed DCs, DCs pulsed with autologous cell lysate, or αCD3/CD28 antibodies from three time points (from **A** and **B**); the experiment using cells from the third time point yielded no virus production.

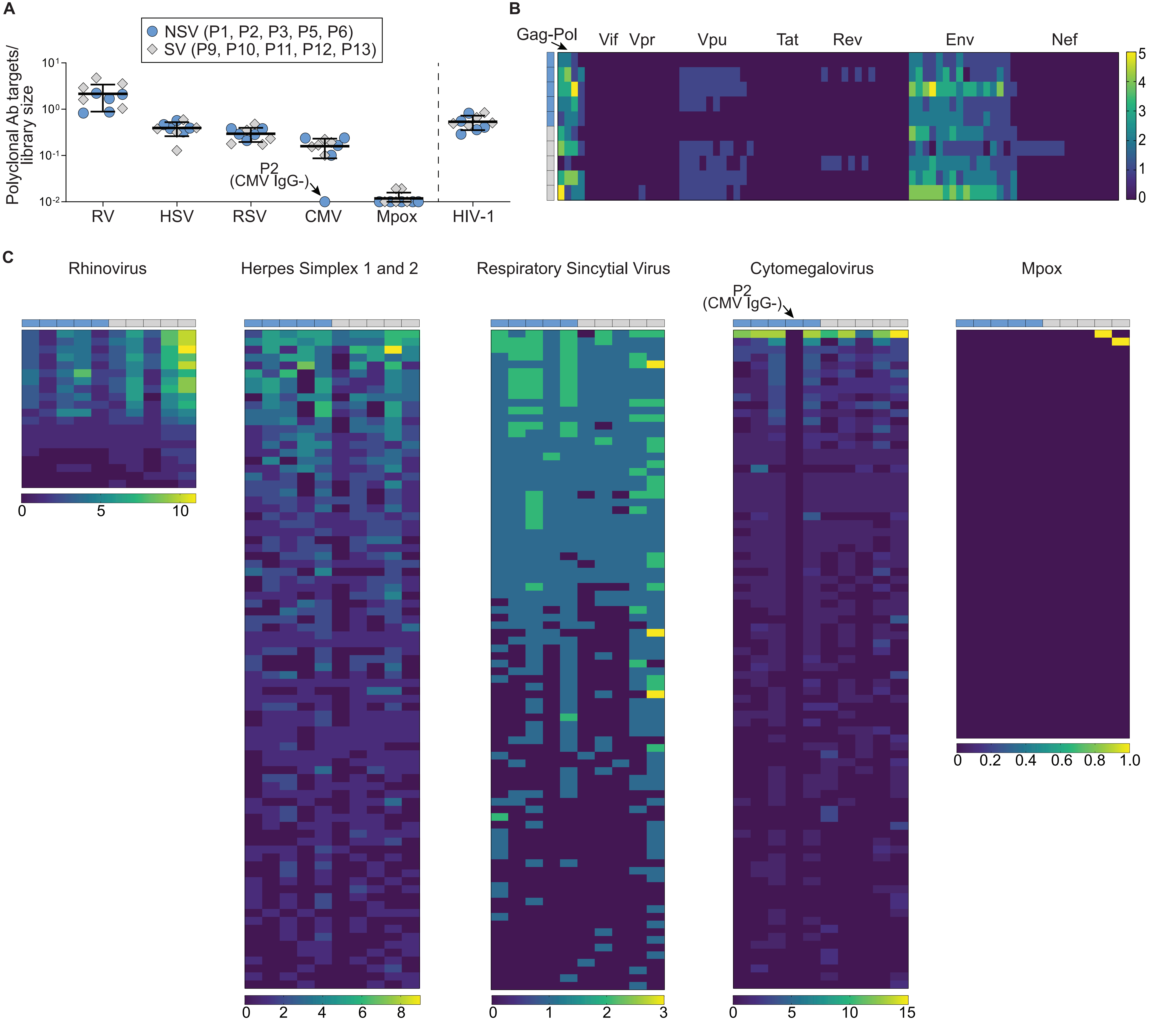


**Supplementary Figure S6. PhIP-seq analysis of polyclonal antibody responses targeting viral proteins.** (**A**) Number of polyclonal antibody responses against viral proteins normalized by protein library size; participants with NSV and suppressed viremia (SV) are indicated by blue circles and grey diamonds, respectively; errors bars show average values and standard deviation. (**B**) Heatmap of antibody responses against HIV-1 with each row representing one participant and each column indicates one protein. The color scale represents the number of antibodies targeting distinct peptides from the same protein. (**C**) Heatmaps as in B, but proteins are ranked by the total sum among participants (highest to lowest); for ease of visualization, heatmaps for HSV, RSV, and CMV show the first 83 proteins.

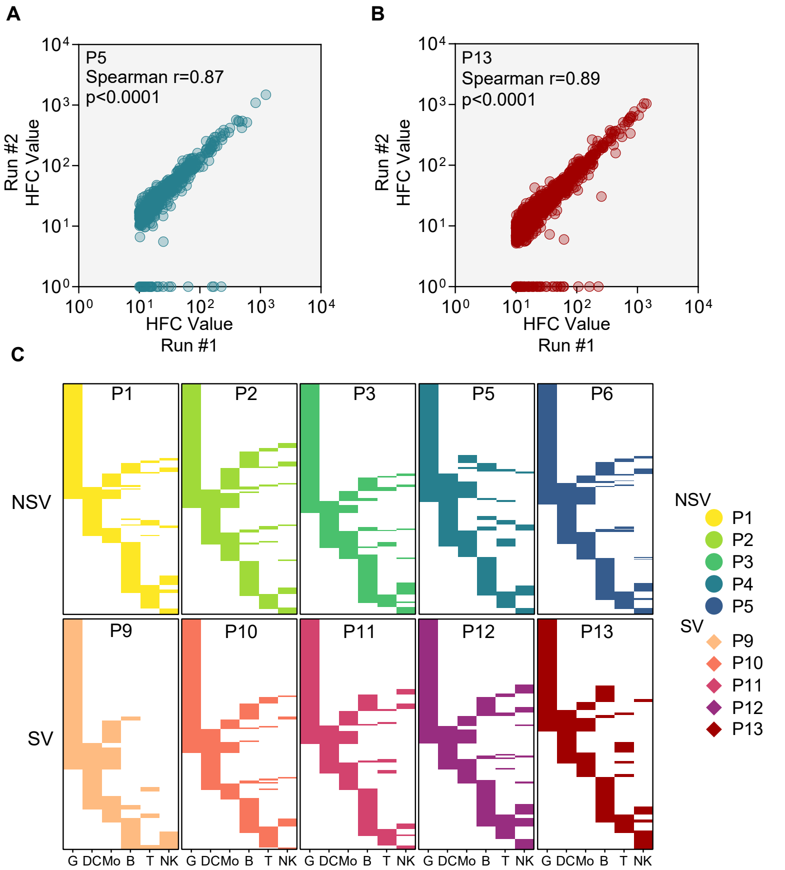


**Supplementary Figure S7. PhIP-seq analysis run validation and autoantibody hit enrichment to immune cell types.** (**A-B**) Correlation plot analysis comparing two PhIP-seq runs for P5 and P13; each dot represents one antibody-reactive peptide form the human peptidome library; axis indicate hit fold changes. (**C**) Immune cell gene enrichment analysis for PhIP-seq hits in all participants. The x axis shows groups of immune cell lineages: G (granulocyte), DC (dendritic cells), Mo (monocytes), B cells, T cells, and natural killer (NK) cells. Each bar represents one autoantibody hit with duplicates removed from the analysis. In the y-axis, each horizontal line represents a peptide that meets the criteria for reactivity (refer to Methods for more information). Lines width is adjusted to a fixed axis height.

| Participant | | Age (Years) | Sex | Race/Etnicity | Current antiretroviral therapy | Years on ART | Last HIV-1 RNA (copies/mL) | Last CD4 count (cells/mm3) |
|---|---|---|---|---|---|---|---|---|
| NSV | P1 | 63 | Male | Black/African American | BIC/FTC/TAF | 9 | 50.8 | 794 |
| | P2 | 60 | Male | Caucasian/White | BIC/FTC/TAF | 26 | 37.8 | 756 |
| | P3 | 58 | Female | Black/African American | BIC/FTC/TAF | 15 | 191 | 761 |
| | P4 | 60 | Male | Caucasian/White | BIC/FTC/TAF/DOR/MVC | 27 | 3400 | 610 |
| | P5 | 55 | Male | Black/African American | BIC/FTC/TAF | *na* | 40.5 | 705 |
| | P6 | 61 | Female | Black/African American | BIC/FTC/TAF | 23 | 129 | 1651 |
| | P7 | 56 | Male | Caucasian/White | ABC/DTG/3TC | 31 | 29.5 | 396 |
| | P8 | 67 | Male | Black/African American | FTC/TAF/RPV | *na* | 31.6 | 273 |
| Suppressed viremia | P9 | 40 | Female | Black/African American | ABC/DTG/3TC | 11 | <20 | 1257 |
| | P10 | 43 | Male | Black/African American | ABC/DTG/3TC | 22 | <20 | 513 |
| | P11 | 62 | Male | Black/African American | BIC/FTC/TAF | 18 | <20 | 475 |
| | P12 | 50 | Male | Black/African American | TDF/FTC/ATZ/r | 11 | <20 | 763 |
| | P13 | 55 | Male | Black/African American | BIC/FTC/TAF | 13 | <20 | 508 |

**Supplementary Table S1. Participant characteristics.** BIC, bictegravir; FTC, emtricitabine; TAF, tenofovir alafenamide; DOR, doravirine; MCV, maraviroc; ABC, abacavir; DTG, dolutegravir; 3TC, lamivudine; RPV, rilpivirine; TDF, tenofovir disoproxil fumarate; ATZ/r, atazanavir boosted with ritonavir.
